# PRMT1 arginine methylation of MCM4 restricts ssDNA gap formation during DNA replication

**DOI:** 10.64898/2026.06.03.729374

**Authors:** Michael W. Ferguson, Cassandra J. Wong, Xiaohan Yang, Edith Cheng, Antonio Mollica, Shahir M. Morcos, Wade H. Dunham, Zhen-Yuan Lin, Eric I. Campos, Anne-Claude Gingras, Grant W. Brown

## Abstract

Replication-associated single-stranded DNA (ssDNA) gaps are increasingly recognized as major sources of genome instability, but how replisome-associated mechanisms suppress their formation remains unclear. Here we identify protein arginine methyltransferase 1 (PRMT1) as a replisome-associated methyltransferase that promotes DNA synthesis and fork integrity. PRMT1 localizes to active forks and its depletion or inhibition slows fork progression in multiple human cell types. We identify MCM4 as a direct PRMT1 substrate and show that methylation-deficient MCM4 causes spontaneous replication stress, elevated RPA, and nuclease-sensitive nascent DNA tracts, indicating persistent daughter-strand gaps. Despite these defects, methylation-deficient MCM4 cells exhibit apparent fork acceleration, consistent with discontinuous DNA synthesis. By contrast, PRMT1 depletion slows forks and selectively reduces fork association of SMC5/6 without broadly disrupting core replisome components. Together, these data support a model in which PRMT1-dependent methylation of MCM4 suppresses gap-prone DNA synthesis, while fork-associated SMC5/6 stabilizes vulnerable gap-containing intermediates.

---

DNA replication ensures the accurate and efficient duplication of genetic material prior to cell division. Genome duplication is mediated by the replisome, a dynamic multiprotein machine that coordinates the synthesis of DNA. Although *in vitro* reconstitutions of the replisome are beginning to achieve the rates of DNA synthesis observed *in vivo*^1^, the full complement of replisome-associated factors and regulatory mechanisms remains incompletely defined. Identifying regulators of DNA synthesis and fork integrity is particularly important because replication forks operate within a crowded and heterogeneous chromosomal environment, where they frequently encounter obstacles that impede, stall, block, or terminate nascent DNA synthesis, a phenomenon broadly referred to as DNA replication stress^2^. Replication stress arises from a variety of endogenous and exogenous sources, and is a hallmark of cancer^3^. Stressors that provoke replication stress include nucleotide depletion, mutations in replication factors, transcription–replication conflicts, and difficult-to-replicate genomic regions. Replication stress challenges cells in two principal ways. First, prolonged periods of replication stress threaten the completion of DNA replication prior to cell division. Second, stressed replication forks accumulate high levels of single-stranded DNA (ssDNA)^4,5^, which is susceptible to mutation, breakage and nucleolytic cleavage, thereby promoting genome instability. In response, the replication stress checkpoint activates ATR kinase to maintain cellular homeostasis, stabilize stressed forks, and prevent aberrant DNA synthesis^6–9^.

Replication stress frequently gives rise to daughter-strand gaps, which are discontinuities in nascent DNA that occur when DNA synthesis becomes uncoupled from fork progression or resumes downstream of replication impediments^10,11^. Daughter-strand gaps can arise following helicase–polymerase uncoupling and downstream repriming^12,13^, delayed post-replicative gap filling^10,14^, and defective Okazaki fragment maturation^15–17^. Although daughter-strand gaps can permit bulk genome duplication to continue, they leave stretches of single-stranded DNA that are vulnerable to mutation, nucleolytic processing, and chromosome breakage if not properly protected and repaired^18–20^. Daughter-strand gaps are therefore increasingly recognized as central intermediates in endogenous replication stress and therapeutic response^21–23^. How replisome-associated mechanisms suppress excessive gap formation and preserve vulnerable gap-containing fork intermediates remains incompletely understood.

DNA unwinding during replication is catalyzed by the CMG helicase, composed of MCM2-7, CDC45, and GINS^24,25^. In addition to its enzymatic role in DNA unwinding, CMG serves as a major organizational platform at replication forks, coordinating DNA synthesis with checkpoint and fork protection factors^26^. Consistent with the importance of proper CMG function, hypomorphic *MCM4* alleles destabilize MCM complex stability, disrupt fork progression, and promote genome instability^27–30^. These observations highlight the role of CMG in limiting abnormal DNA synthesis.

Protein arginine methyltransferases (PRMTs) catalyze the methylation of arginine residues. Mammalian cells express nine PRMTs comprising three catalytic classes. Type I PRMTs catalyze asymmetric dimethylation, type II PRMTs catalyze symmetric dimethylation, and type III PRMTs catalyze monomethylation^31,32^. Arginine methylation is a pervasive post-translational modification, with proteome-wide prevalence comparable to phosphorylation^33^, influencing protein localization^34^, enzyme activity^35^, protein-protein interactions^36,37^, and cell signalling^32^. Although glycine- and arginine-rich (GAR) motifs are common arginine methylation sites, increasing evidence indicates a broader substrate specificity for PRMTs^32,33^. PRMT1 is the predominant type I PRMT, accounts for ∼85% of all cellular arginine methylation^38,39^, and is essential in human cells^40^ and mouse embryos^41^. PRMT1 is well established as a regulator of gene expression^42–45^ and genome maintenance^35,46–48^, and has also been linked to replication stress responses through regulation of ATR abundance^49,50^. Recently, PRMT1 has been linked to reduced replication fork progression in mouse hematopoietic stem cells^51^, suggesting that its functions may extend beyond transcription and DNA repair. Whether PRMT1 acts directly at active replisomes, which replication proteins are functionally methylated by PRMT1, and how arginine methylation regulates fork mechanics remain unknown.

Here, we identify PRMT1 as a previously unappreciated replisome-associated regulator of DNA replication. We show that PRMT1 localizes to active replisomes and that chemical inhibition or depletion of PRMT1 impairs DNA synthesis and alters replication fork composition. We identify MCM4 as a direct PRMT1 substrate and show that arginine methylation of MCM4 suppresses aberrant fork progression and the formation of replication-associated daughter-strand ssDNA gaps. We further uncover a role for the SMC5/6 complex in stabilizing vulnerable gap-containing fork intermediates. Together, our findings establish PRMT1-dependent arginine methylation as a replisome-centered mechanism that promotes continuous DNA synthesis, preserves fork integrity, and safeguards genome stability.

## Results

### PRMT1 is enriched at DNA replication forks

To identify novel DNA replication factors, we performed isolation of proteins on nascent DNA (iPOND) mass spectrometry (MS)^52^ in HEK293 cells. After briefly pulse-labelling with 5-ethynyl-2′-deoxyuridine (EdU), cells were chemically cross-linked, and newly synthesized DNA was biotinylated using click chemistry. Proteins proximal to nascent DNA were captured by streptavidin pulldown and identified using mass spectrometry. Proteins identified in the EdU pulse with a significance analysis of interactome (SAINT) score^53^ ≥ 0.95 and at least 6 spectral counts across two replicates were considered to be robust nascent DNA interactors, of which there were 177 proteins. We then filtered for those proteins showing a ≥ 2-fold enrichment in spectral counts in the EdU pulse relative to a thymidine chase to identify 82 proteins that localize to active replisomes. The 82 proteins at DNA replication forks are presented as nodes of a STRING network in Figure 1A, with edges marking high-confidence protein-protein interactions^54^. As expected, we identified proteins with known roles in DNA replication including the MCM helicase (MCM2-6), clamp and clamp loader (PCNA, RFC1-5), and polymerase subunits (POLA1, PRIM2, POLD1, POLD3, POLE). Furthermore, we identified proteins with chromatin-associated functions that were enriched at replisomes, including members of the NuRD complex (MTA1, MTA2, HDAC1, HDAC2) and SWI/SNF chromatin remodelers (SMARCA1, SMARCA5). Gene ontology analysis of the corresponding 82 genes revealed strong enrichments for the regulation of DNA replication, DNA damage response, and chromatin remodeling (Figure 1B).

**Figure 1.**
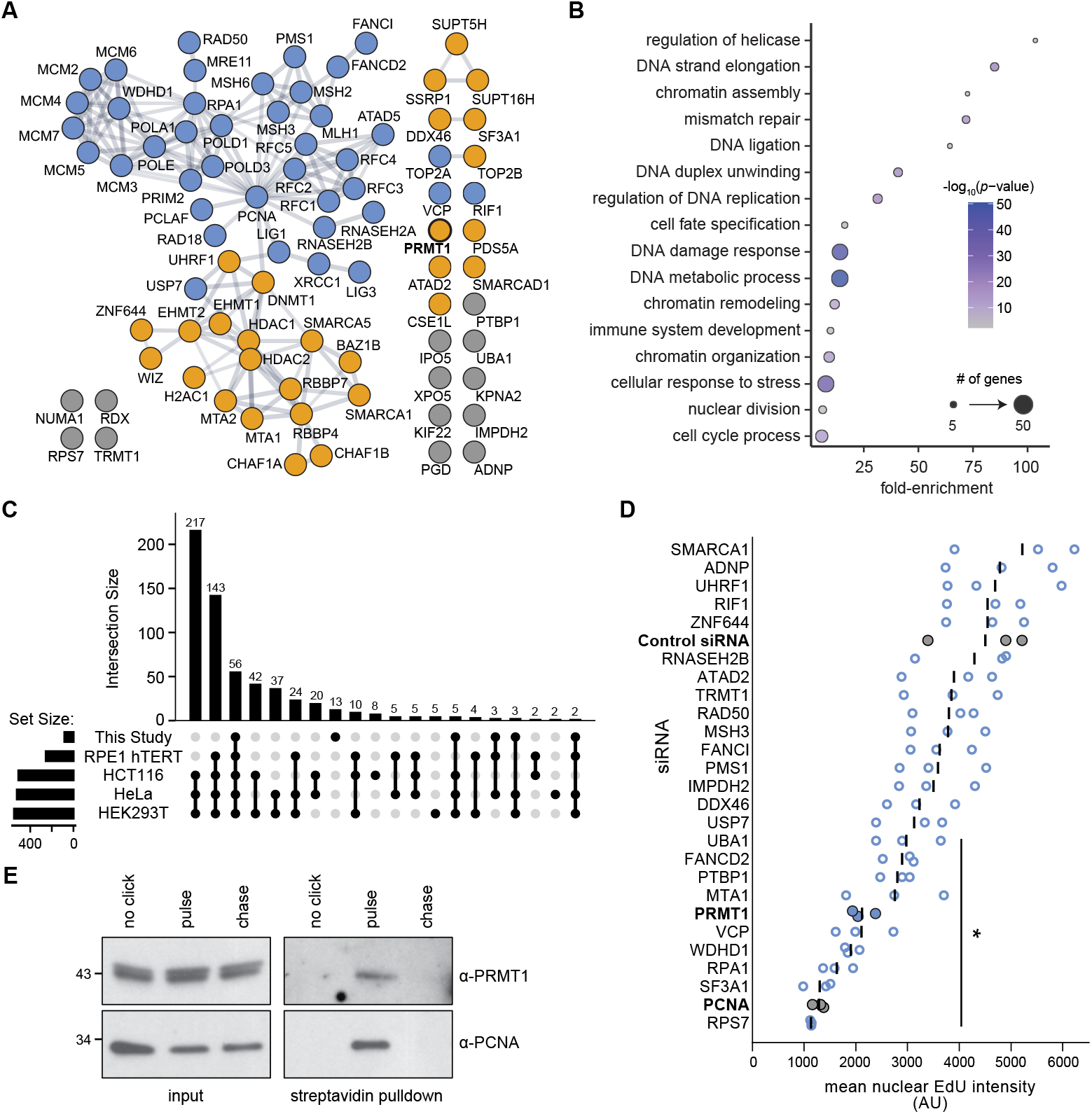
PRMT1 is enriched at DNA replication forks. (A) STRING network of the 82 proteins enriched at replisomes in iPOND-mass spectrometry. Edges represent known high-confidence protein-protein interactions and edge thickness is weighted based on the confidence of the interaction. Manually curated gene functions are indicated by node colour: blue, DNA replication; orange, chromatin-associated; grey, other. *n*=2 biological replicates. (B) Gene ontology (GO) biological process analysis for the 82 high-confidence proteins that localize to active replisomes. The fold-enrichment for each GO term is indicated, colours indicate corrected *p*-values, and the size of the circles corresponds to the number of genes annotated to each GO term. (C) UpSet plot showing the intersections of our study and iPOND-mass spectrometry datasets from Wessel *et* al^55^. Filled circles connected by vertical lines denote the datasets represented by each intersection. Vertical bars indicate the number of proteins in each intersection, and horizontal bars indicate the total number of proteins in each dataset. Intersections are ordered by decreasing size and proteins unique to each dataset are indicated by the absence of vertical bars. (D) EdU incorporation following siRNA knockdown of the indicated genes. Following knockdown, cells were pulse-labelled with EdU. Nuclear EdU intensity was measured by fluorescence microscopy and is plotted in arbitrary units (AU). Circles represent the mean nuclear EdU intensity of each replicate, and the black bars indicate the average of the replicates. Knockdowns with statistically supported decreases in nuclear EdU intensity relative to the control siRNA (*p* < 0.05; one-tailed Student’s t-test) are indicated *. siPCNA served as the positive control. *n*=3 biological replicates. (E) HEK293T cells were subjected to the iPOND workflow, and the input extracts and streptavidin pulldowns were immunoblotted for the presence of PRMT1, PCNA (DNA replication fork control), and histone H3 (chromatin control). The control without covalent attachment of streptavidin to EdU (no click), the EdU pulse, and the thymidine chase are shown. The positions of molecular weight standards, in kDa, are indicated to the left.

We compared our data to a previous study where iPOND was coupled to stable isotope labeling by amino acids in cell culture (SILAC) mass spectrometry to identify proteins enriched on nascent DNA in four cell lines^55^. 56 of our hits (68.2%) were identified in all four cell lines investigated by Wessel *et al*., while 13 (15.9%) of the proteins enriched at replication forks were unique to our study (Figure 1C). Therefore, our iPOND-MS dataset largely recaptures the protein environment at replisomes and successfully identifies DNA replication fork proteins.

To identify proteins with unrecognized roles in DNA replication, we performed a secondary screen of 25 genes to assess their roles in promoting DNA synthesis (Figure 1D). We knocked down each gene with a pool of siRNAs and measured EdU incorporation after 48 hours. Interestingly, 10 different knockdowns led to a statistically supported decrease in the mean nuclear EdU intensity of cells, suggesting that our iPOND experiment identified factors that promote DNA synthesis. Of particular interest, we identified the arginine methyltransferase PRMT1 in our secondary screen. A role for PRMT1 in promoting DNA synthesis through physical association with replication forks has not been described. To confirm the PRMT1 enrichment at replication forks detected by iPOND-MS, we performed iPOND-western blotting in HEK293 cells (Figure 1E). Consistent with our iPOND-MS data, and with that of Wessel *et al*^55^, PRMT1 associated with nascent DNA (pulse) but was not detected on post-replicative DNA (chase). Together, our data demonstrate that PRMT1 is enriched at active replisomes.

### PRMT1 promotes DNA synthesis

We next tested whether PRMT1 promotes DNA synthesis in human cells using two orthogonal techniques. *PRMT1* is an essential gene^40^ and therefore is not amenable to stable gene knockout. We generated a doxycycline-regulated CRISPR-interference (CRISPRi) system^56^ in A375 cells to conditionally deplete *PRMT1* expression. PRMT1 was robustly depleted by two different guide RNAs following doxycycline treatment (Figure 2A), as was global asymmetric arginine dimethylation (Figure S1A). We performed EdU flow cytometry to measure DNA synthesis following PRMT1 depletion and observed a marked decrease in the amount of EdU incorporated into the nascent DNA of S-phase cells following a 3- or 5-day depletion of PRMT1 (Figures 2B and S1B). Next, we measured DNA replication fork rates directly using DNA combing^57,58^. Consistent with the decrease in EdU incorporation, we measured a 20-23% decrease in fork rates when PRMT1 was depleted by CRISPRi (Figure 2C). These results were recapitulated in U2OS cells where fork rate decreased by 27-33% following a 2-day siRNA knockdown of *PRMT1* (Figures S1C and S1D). We infer that in addition to being enriched at DNA replication forks, PRMT1 is important for promoting DNA synthesis in human cells.

**Figure 2.**
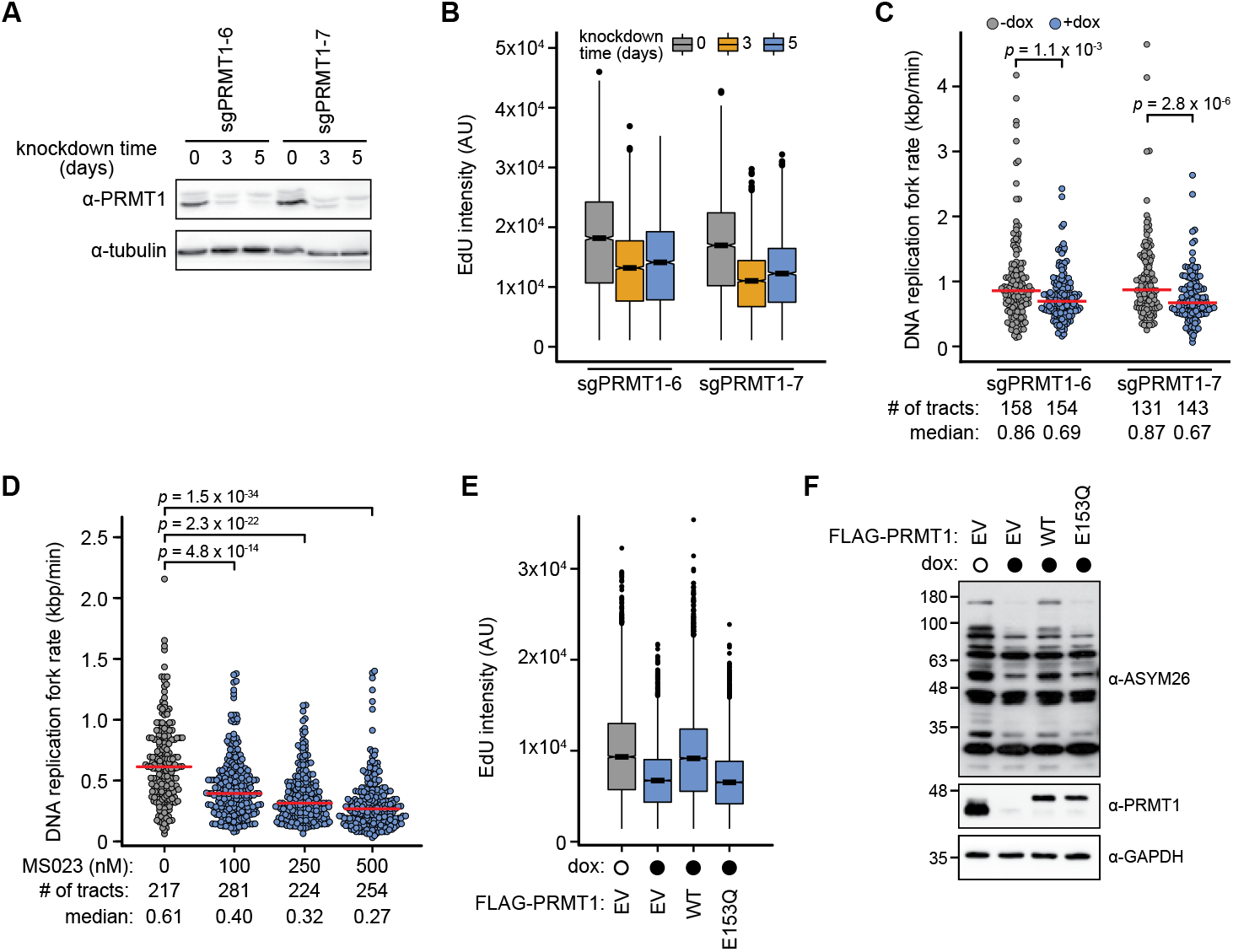
PRMT1 promotes DNA synthesis. (A) PRMT1 depletion in CRISPRi lines. CRISPRi A375 cells carrying doxycycline-inducible sgRNAs for *PRMT1* were sampled at the indicated times after the addition of doxycycline and immunoblotted to detect PRMT1 and tubulin. The positions of molecular weight standards, in kDa, are indicated to the left. (B) EdU incorporation in *PRMT1* CRISPRi cells. CRISPRi cells carrying doxycycline-inducible sgRNAs for *PRMT1* were pulse-labelled with EdU at the indicated times after the addition of doxycycline. EdU intensity per cell was measured by flow cytometry and is plotted in arbitrary units as box plots, with horizontal bars indicating the medians. Boxes span the first through third quartiles, whiskers extend to the last data points within 1.5 times the interquartile range, and outliers are plotted as circles. A minimum of 11,000 cells were measured per sample. *n*=2 biological replicates (replicate 2, Figure S1B). (C) DNA combing analysis of *PRMT1* CRISPRi cells. PRMT1 was depleted for 5 days by expressing the indicated sgRNAs (+dox), and cells were sequentially pulse-labelled with CldU and IdU, followed by DNA fibre isolation and molecular combing. The IdU tract lengths were measured and plotted as replication fork rates. Parallel cultures without PRMT1 depletion are shown for comparison (-dox). Medians are indicated by horizontal red bars and *p*-values were calculated with a two-sided Mann-Whitney *U* test. (D) DNA combing analysis of HEK293T cells following type I PRMT inhibition. Cells were treated with MS023 at the indicated concentrations for 48 hours, and sequentially pulse-labelled with CldU and IdU, followed by DNA fibre isolation and molecular combing. The IdU tract lengths were measured and plotted as replication fork rates. Parallel cultures without PRMT1 depletion are shown for comparison (-dox). Medians are indicated by horizontal red bars and *p*-values were calculated with a two-sided Mann-Whitney *U* test. (E) Rescue of EdU incorporation in *PRMT1* CRISPRi cells. CRISPRi sgPRMT1-7 cells were treated with doxycycline for 5 days (closed circles) to deplete PRMT1, prior to transfection with the empty vector (EV), *FLAG-PRMT1* (WT), or *FLAG-PRMT1*_*E153Q*_ (catalytic-dead; E153Q) prior to pulse-labelling with EdU, EdU intensity per cell was measured by flow cytometry and is plotted in arbitrary units as box plots, with horizontal bars indicating the medians. Boxes span the first through third quartiles, whiskers extend to the last data points within 1.5 times the interquartile range, and outliers are plotted as circles. A parallel culture without PRMT1 depletion is shown for comparison (open circle). *n*=2 biological replicates (replicate 2, Figure S1H). (F) Whole cell lysates from the samples in (E) were immunoblotted to detect asymmetric arginine dimethylation (α-ASYM26), PRMT1, and GAPDH. The positions of molecular weight standards, in kDa, are indicated to the left.

Another arginine methyltransferase, CARM1 (PRMT4), localizes to replication forks and plays a non-enzymatic role in regulating fork speed via its interaction with PARP1^59^. Thus, we tested whether the enzymatic activity of PRMT1 was required to promote DNA synthesis. We first used the type I PRMT inhibitor MS023 to disrupt binding of PRMT1 to its arginine substrates^60^. MS023 treatment of HEK293T cells led to a substantial decrease in replication fork rate as measured by DNA combing (Figure 2D) and a decrease in global asymmetric arginine dimethylation (Figure S1E). U2OS and A375 cells treated with MS023 also displayed decreased EdU incorporation into nascent DNA (Figures S1F and S1G), indicating that arginine substrate binding by a type I PRMT is required to maintain fork speed. Since MS023 is not specific for PRMT1, we performed a rescue experiment by constitutively expressing either wild-type or enzymatically inactive PRMT1 (E153Q)^61^ while depleting endogenous PRMT1. Expression of wild-type but not mutant PRMT1 rescued the DNA synthesis defect observed when endogenous PRMT1 was depleted (Figures 2E and S1H). As expected, wild-type but not mutant PRMT1 also rescued global asymmetric arginine dimethylation (Figures 2F and S1I). Thus, we conclude that the arginine methyltransferase activity of PRMT1 is required to promote DNA synthesis.

### PRMT1 depletion causes replication stress and DNA damage

Given that PRMT1 depletion results in slowing of DNA replication, we tested whether the abnormal replication kinetics resulted in replication stress and DNA damage. DNA replication stress is often indicated by the accumulation of ssDNA^2,4,62^. We measured the number of RPA foci, as a proxy for ssDNA, in S-phase cells following *PRMT1* CRISPRi (Figure 3A). We did not detect an increase in RPA foci when PRMT1 was depleted. However, when we added exogenous replication stress in the form of the DNA polymerase inhibitor aphidicolin, a dose-dependent increase in RPA foci was evident in the PRMT1-depleted cells. Consistent with the presence of RPA foci, we found that RPA32 Ser 33 phosphorylation increased when *PRMT1* CRISPRi was combined with aphidicolin (Figure 3B, 0.3h samples), indicating activation of the ATR checkpoint kinase. We also detected a modest increase in RPA phosphorylation upon *PRMT1* CRISPRi in the absence of aphidicolin, suggesting that PRMT1 depletion alone causes sufficient DNA replication stress to activate the ATR checkpoint response.

**Figure 3.**
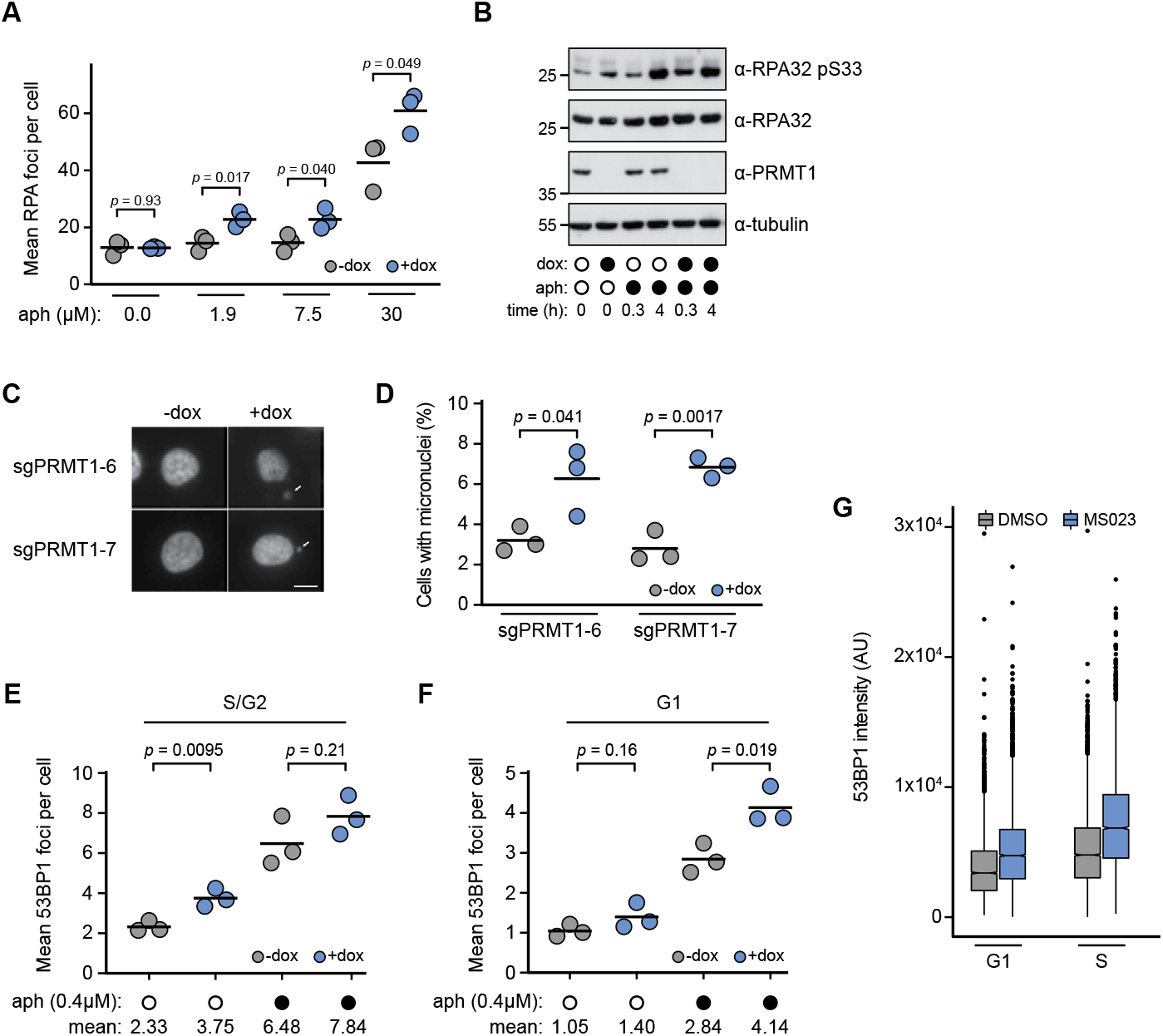
PRMT1 depletion induces replication stress and DNA damage. (A) RPA foci measured by immunofluorescence microscopy in CRISPRi sgPRMT1-6 cells following a 5-day PRMT1 depletion (+dox), a 30-minute aphidicolin treatment (aph) at the indicated concentration, and EdU pulse-labelling. EdU-positive (S-phase) cells were analyzed. The mean number of RPA foci per cell in each replicate is plotted, with horizontal bars indicating the means of the replicates. *n*=3 biological replicates. (B) RPA phosphorylation in *PRMT1* CRISPRi cells. Whole cell lysates of *PRMT1* CRISPRi cells were prepared following a 5-day PRMT1 depletion (+dox), and treatment with aphidicolin (aph) for the indicated times, and immunoblotted. The immunoblots were probed for the middle subunit of RPA (RPA32), phosphorylated RPA32 (RPA32 pS33), PRMT1, and tubulin. The positions of molecular weight standards, in kDa, are indicated to the left. (C) Micronuclei in *PRMT1* CRISPRi cells. Representative micrographs of *PRMT1* CRISPRi cells following a 5-day PRMT1 depletion (+dox). Cells were stained with DAPI prior to fluorescence microscopy. Micronuclei are indicated by the arrows, and untreated cells (-dox) are shown for comparison. Scale bar = 10 µm. (D) The percent of PRMT1-depleted (+dox) and untreated (-dox) cells with micronuclei is plotted for each replicate. The means are indicated by the horizontal bars, and *p*-values were calculated using a two-tailed Student’s t-test. *n*=3 biological replicates. (E) 53BP1 foci were measured by immunofluorescence microscopy in ^2^ CRISPRi sgPRMT1-6 cells following a 5-day PRMT1 depletion (+dox) and a 16 h aphidicolin treatment (aph). Cells were treated with anti-cyclin A antibody to identify cells in S and G2 phases (S/G2). The mean number of 53BP1 foci per S/G2 cell is plotted for each replicate, means of the replicates are indicated by the horizontal bars, and *p*-values were calculated using a two-tailed Student’s t-test. *n*=3 biological replicates. (F) 53BP1 foci were measured as in (E), for G1-phase cells. (G) Quantification of chromatin-bound 53BP1 following PRMT inhibition. U2OS cells were treated with MS023 for 3 days prior to pulse-labelling with EdU. Cells were extracted to remove soluble proteins, stained with DAPI, and subjected to flow cytometry. 53BP1 intensity was measured for each cell in G1 phase or S phase and plotted in arbitrary units as box plots, with horizontal bars indicating the medians. Boxes span the first through third quartiles, whiskers extend to the last data points within 1.5 times the interquartile range, and outliers are plotted as circles. A minimum of 11,000 cells were measured per sample. *n*=2 biological replicates (replicate 2, Figure S2A).

Unresolved replication stress can manifest in micronuclei formation during mitosis^63^. We found that PRMT1 depletion increased the number of micronuclei by 2.0-to 2.4-fold (Figures 3C and 3D). Next, we quantified the number of 53BP1 foci per cell, a marker of DNA double-strand breaks^64,65^, in the presence and absence of aphidicolin-induced replication stress (Figures 3E and 3F). When PRMT1 was depleted, we observed a modest increase in 53BP1 foci per cell during S/G2, suggesting that DNA breaks are elevated when replication occurs in the absence of PRMT1 (Figure 3E). The accumulation of breaks in S/G2 was not exacerbated by ectopic replication stress (Figure 3E, 0.4 µM aph). When we examined G1 cells, we found that the level of breaks was similar in the presence and absence of PRMT1 (Figure 3F). However, the addition of aphidicolin resulted in more 53BP1 foci following PRMT1 depletion. Treating U2OS cells with MS023 caused elevated levels of chromatin-bound 53BP1 in G1 and S-phase cells (Figures 3G and S2A). Together our data are consistent with PRMT1 depletion causing under-replication that results in chromosome breakage and micronuclei formation in mitosis^66,67^ and persistent double-strand DNA breaks in the subsequent G1 phase^68–70^.

### MCM4 is asymmetrically arginine dimethylated by PRMT1

Since PRMT1 is enriched at replication forks, and the arginine methyltransferase activity of PRMT1 is necessary for normal replication fork progression, we reasoned that PRMT1 could arginine methylate a replication protein to promote proper DNA synthesis. We compared our iPOND-MS dataset to a quantitative methylproteomics analysis that measured changes in monomethyl arginine abundance in response to *PRMT1* knockdown^33^ (Figure 4A). We found that 14 proteins were enriched at active DNA replication forks and arginine monomethylated in a PRMT1-dependent manner (Figure 4A), including two known substrates of PRMT1, MRE11 and SUPT5H^35,71^. Interestingly, the methylproteomics analysis detected substantial arginine monomethylation of the MCM heterohexamer, a component of the CMG helicase, and the methylation of MCM6 and MCM7 was PRMT1-dependent^33^. Furthermore, arginine methylation of MCM2 and MCM4 was detected in a parallel proteomics analysis^72^, and MCM2, MCM6, MCM7 and PRMT1 form a stable protein complex^73^. Therefore, we tested whether members of the MCM complex were asymmetrically arginine dimethylated. We expressed and immunoprecipitated each MCM subunit and assessed arginine methylation status by immunoblotting. We detected robust asymmetric arginine dimethylation of MCM4 tagged either with V5 or eGFP, but not other MCM subunits (Figures 4B and S3A). Although an arginine dimethylated species was present in the MCM2-V5 pulldown, it co-migrated with MCM4 and was not evident in MCM2-eGFP pulldowns. We conclude that MCM4 is asymmetrically arginine dimethylated and is therefore a type I arginine methyltransferase target.

**Figure 4.**
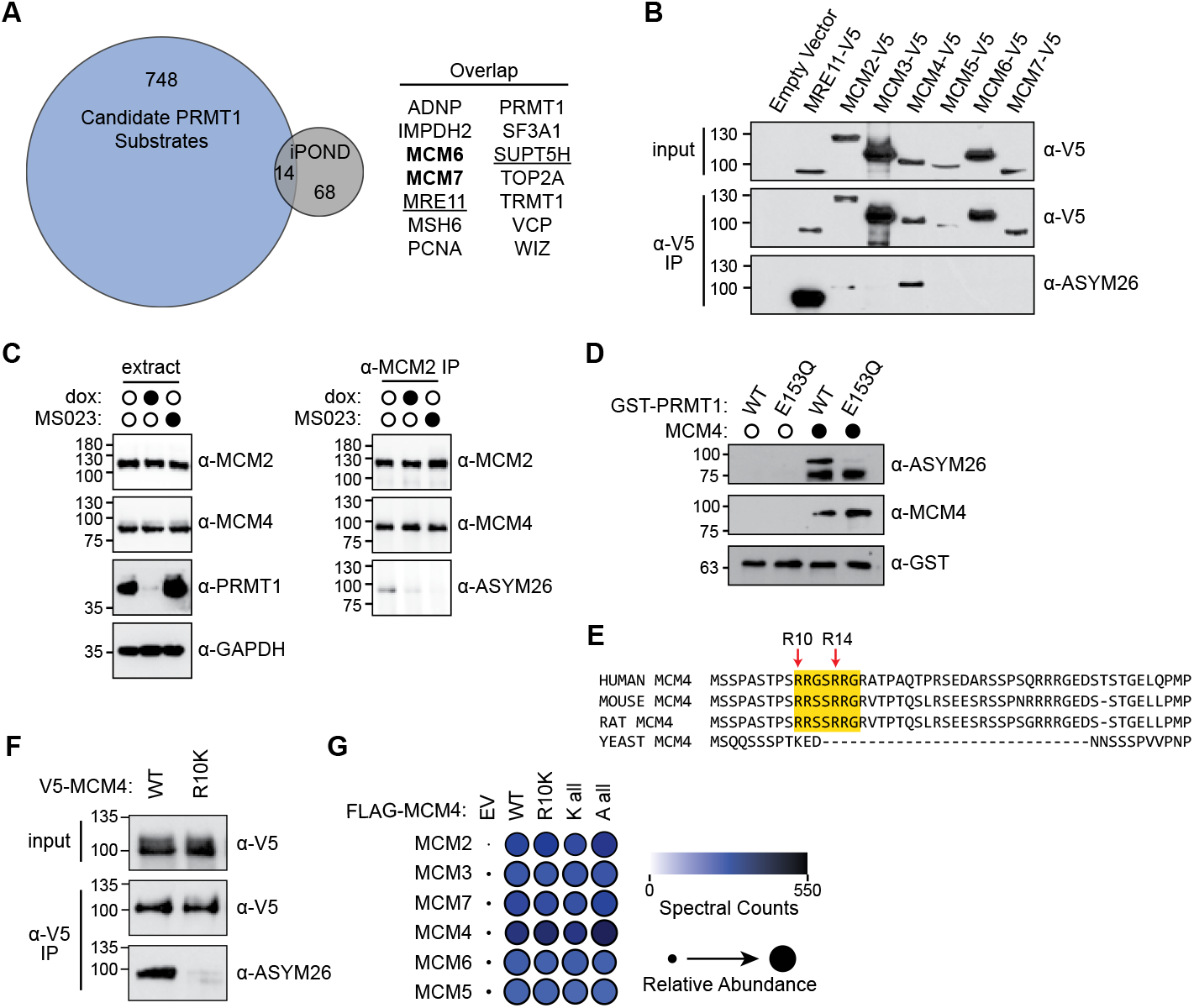
PRMT1 asymmetrically arginine dimethylates MCM4. (A) Venn diagram of the overlap between the Larsen et al methylproteomics study33 and replisome-enriched proteins from Figure 1A. Proteins whose monomethylation abundance changed upon PRMT1 depletion are in the blue circle, and proteins localized to DNA replication forks are in the grey circle. The 14 proteins present in both datasets are shown to the right, with known substrates of PRMT1 underlined and MCM subunits in bold. (B) Asymmetric arginine dimethylation of MCM subunits analyzed by immunoblotting. HEK293T cells were transfected to express the indicated V5-tagged MCM proteins or MRE11-V5, nuclear extracts were prepared (input), and immunoprecipitated with an anti-V5 antibody (α -V5 IP). Immunoblots were probed to detect the V5-tagged proteins and asymmetric arginine dimethylation (α-ASYM26). The positions of molecular weight standards, in kDa, are indicated to the left. (C) Asymmetric arginine dimethylation of the MCM complex analyzed following PRMT1 depletion or inhibition. Whole-cell (left) and nuclear (right) extracts were prepared from CRISPRi sgPRMT1-6 cells following a 5-day PRMT1 depletion (+dox) or 2 days of PRMT1 inhibition with 1 µM MS023. An α -MCM2 antibody was used to precipitate the MCM complex from the nuclear extracts (α -MCM2 IP), and the whole-cell extracts and IPs were immunoblotted. The immunoblots were probed to detect MCM2, MCM4, PRMT1, GAPDH, and asymmetric arginine demethylation GAPDH, and asymmetric arginine dimethylation (α-ASYM26). The positions of molecular weight standards, in kDa, are indicated to the left. (D) Methylation of recombinant MCM4 with purified GST-PRMT1. MCM4 was incubated with wild-type (WT) or catalytically inactive (E153Q) GST-PRMT1 for 3 hours at 30°C. The samples were immunoblotted to detect MCM4, GST-PRMT1, and asymmetric arginine dimethylation (α-ASYM26). The positions of molecular weight standards, in kDa, are indicated to the left. (E) Multiple sequence alignment of *Homo sapiens, Mus musculus, Rattus norvegicus*, and *Saccharomyces cerevisiae* N-terminal MCM4 protein sequences. The putative GAR motif is highlighted with a yellow box, and the red arrows mark arginine residues detected as dimethylated by mass spectrometry^72^. (F) Asymmetric arginine dimethylation of MCM4 variants. HEK293T cells were transfected to express V5-tagged MCM4 (WT) or MCM4_R10K_ (R10K). Nuclear extracts were prepared (input), precipitated with anti-V5 antibody (α-V5 IP), and immunoblotted to detect the V5-MCM4 proteins and asymmetric arginine dimethylation (α-ASYM26). The positions of molecular weight standards, in kDa, are indicated to the left. (G) Assessment of MCM4 variant interactions with the MCM2-7 complex. FLAG-tagged wild-type MCM4 (WT), MCM4_R10K_ (R10K), MCM4_Kall_ (K all), or MCM4_Aall_ (A all), or the empty vector (EV) were integrated in HEK293 Flp-In T-Rex cells, and expression was induced for 16 hours. The MCM4-FLAG proteins were immunoprecipitated (α-FLAG IP) and analyzed by mass spectrometry. The number of spectral counts and relative abundance of each MCM2-7 complex member is plotted.

We next tested if the asymmetric arginine dimethylation of MCM4 was PRMT1-dependent. We depleted PRMT1 with CRISPRi or inhibited PRMT1 with MS023, immunoprecipitated the MCM complex from nuclear extracts using an anti-MCM2 antibody, and probed immuoblots for asymmetric arginine dimethylation (Figure 4C). We detected asymmetric arginine dimethylation of a ∼90 kDa protein that co-migrated with MCM4 in immunoprecipitates from cells expressing PRMT1. When PRMT1 was depleted or chemically inhibited, arginine dimethylation of the ∼90 kDa protein decreased substantially. We then performed *in vitro* methylation with purified recombinant proteins (Figure S3B) and detected robust asymmetric arginine dimethylation of MCM4 upon incubation with wild type, but not catalytically inactive, PRMT1 (Figures 4D and S3C). We conclude that MCM4 is a PRMT1 substrate both *in vivo* and *in vitro*.

Mass spectrometric analyses have identified two MCM4 residues that are arginine dimethylated, Arg10 and Arg14^72^. We noted that this region of MCM4 resembles the GAR motif, the canonical methylation site for PRMTs^32^, and is highly conserved across metazoan MCM4 proteins (Figure 4E). We directly tested Arg10 by generating an MCM4 R10K variant, overexpressing MCM4_R10K_, and measuring asymmetric arginine dimethylation (Figure 4F). Strikingly, MCM4_R10K_ had substantially decreased arginine methylation when compared to wild-type MCM4. We expressed three additional *MCM4* GAR motif variants, MCM4_R14K_, MCM4_Kall_ (R10K/R11K/R14K/R15K/R17K), and MCM4_Aall_ (R10A/R11A/R14A/R15A/R17A) (Figure S3D), and found in each case arginine methylation was largely eliminated (Figure S3E). Therefore, we conclude that arginine methylation of MCM4 depends very strongly on Arg10 and Arg14, and likely occurs within the GAR motif.

### MCM4 asymmetric arginine dimethylation does not regulate MCM complex assembly

MCM4 must associate with the 5 other members of the MCM heterohexamer in order to form an active DNA helicase24. We tested whether MCM4 arginine methylation was important for MCM complex formation. Each MCM4 methylation-deficient variant co-immunoprecipitated MCM2 at similar levels to wildtype MCM4 (Figure S3E). Furthermore, affinity purification-mass spectrometric (AP-MS) analysis of FLAG-tagged MCM4 immunoprecipitates showed similar levels of all MCM heterohexamer members with wild-type MCM4, MCM4_R10K_, MCM4_Kall_, and MCM4_Aall_ (Figure 4G). Thus, we conclude that arginine methylation of MCM4 is not required for MCM complex assembly or stability

### Methylation-deficient MCM4 causes DNA replication stress

Mutations in replication factors are well established sources of DNA replication stress^28,29,74^. Therefore, we edited the *MCM4* locus in HAP1 cells to generate clonal lines carrying V5-tagged methylation-deficient *MCM4* alleles (Figure S4A) and determined whether they exhibit signs of replication stress. All knock-in lines expressed MCM4 at similar levels (Figures 5A), and the wild-type V5-MCM4 knock-in remained robustly asymmetrically arginine dimethylated. Arginine methylation was greatly reduced for MCM4_R10K_ and not detected for MCM4_Kall_ and MCM4_Aall_ (Figure 5A).

**Figure 5.**
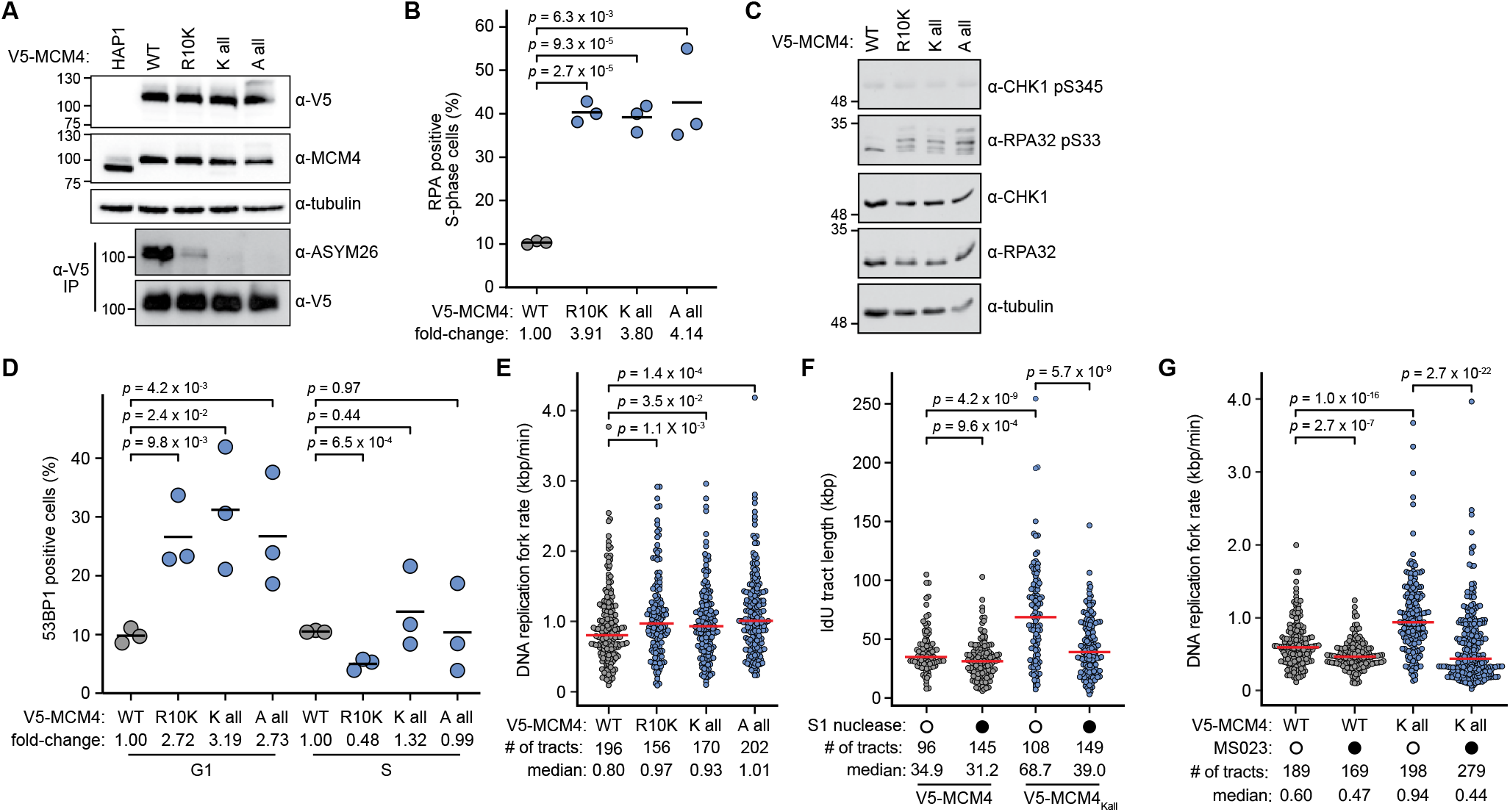
Methylation-deficient MCM4 causes replication stress. (A) *MCM4* knock-in lines analyzed by immunoblotting. Lysates prepared from the indicated HAP1 knock-in lines carrying the indicated V5-tagged *MCM4* variants at the *MCM4* locus were immunoblotted and probed for V5, MCM4, and tubulin. Lysates from the polyclonal parent lines were immunoprecipitated with α-V5 and immunoblotted to detect asymmetric arginine dimethylation with ASYM26. The positions of molecular weight standards, in kDa, are indicated to the left. (B) Quantification of chromatin-bound RPA during S phase in *MCM4* variant knock-in cells. Cells were pulse-labelled with EdU, extracted to remove soluble proteins, treated with α-RPA32 antibody, stained with DAPI, and subjected to flow cytometry. The percent of RPA positive S-phase cells is plotted for each replicate. The means are indicated by the horizontal bars, and *p*-values were calculated using a two-tailed Student’s t-test. *n*=3 biological replicates, and a minimum of 2,750 S-phase cells were analyzed per sample. (C) Analysis of CHK1 and RPA32 phosphorylation in *MCM4* variant knock-in cells. Extracts of the indicated knock-in lines were immunoblotted to detect CHK1 phospho-serine345, RPA32 phospho-serine33, total CHK1, total RPA32, and tubulin. The positions of molecular weight standards, in kDa, are indicated to the left. (D) Quantification of chromatin-bound 53BP1 in *MCM4* variant knock-in cells. The indicated knock-in lines were pulse-labelled with EdU, extracted to remove soluble proteins, stained with DAPI, and subjected to flow cytometry. The percent of G1- or S-phase 53BP1 positive cells is plotted for each replicate. The means are indicated by the horizontal bars, and *p*-values were calculated using a two-tailed Student’s t-test. *n*=3 biological replicates, and a minimum of 7,000 cells were analyzed per sample. (E) DNA combing analysis of *MCM4* knock-in lines. Cells were sequentially pulse-labelled with CldU and IdU, followed by DNA fibre isolation and molecular combing. The IdU tract lengths were measured and are plotted as replication fork rates. Medians are indicated by horizontal red bars and *p*-values were calculated with a two-sided Mann-Whitney *U* test. (F) Nuclease sensitivity of replication tracts. *MCM4* and *MCM4*_*Kall*_ knock-in cells were sequentially pulse-labelled with CldU and IdU, followed by DNA fibre preparation, and S1 nuclease treatment as indicated. After molecular combing the IdU tract lengths were measured and are plotted. Medians are indicated by horizontal red bars and *p*-values were calculated with a two-sided Mann-Whitney *U* test. *n*=2 biological replicates (replicate 2, Figure S4C). (G) DNA replication fork rate analysis of MCM4 knock-in lines following PRMT inhibition. *MCM4* and *MCM4*_*Kall*_ knock-in cells were treated with 2 µM MS023 for 3 days, followed by sequential pulse-labelling with CldU and IdU, DNA fibre isolation, and molecular combing. The IdU tract lengths were measured and are plotted as replication fork rates. Medians are indicated by horizontal red bars and *p*-values were calculated with a two-sided Mann-Whitney *U* test. *n*=2 biological replicates (replicate 2, Figure S4D).

To determine whether expressing methylation-deficient MCM4 caused replication stress, we first quantified the levels of chromatin-bound RPA during S-phase (Figure 5B). Methylation-deficient MCM4 cells displayed a dramatic 4-fold increase in cells with RPA chromatin binding. Loss of MCM4 arginine methylation also increased RPA32 Ser33 phosphorylation, indicating activation of the replication stress checkpoint kinase ATR (Figure 5C). Surprisingly, phosphorylation of the ATR target CHK1 was not detected in our methylation-deficient MCM4 lines (Figure 5C and S4B). Addition of etoposide to induce DNA replication stress resulted in phosphorylation of both RPA32 and CHK1 in methylation-deficient MCM4 cells (Figure S4B), indicating that the absence of CHK1 phosphorylation is specific to MCM4-induced replication stress. Together, our data demonstrate that the loss of MCM4 asymmetric arginine dimethylation induces spontaneous replication stress, activates ATR signaling, yet does not support CHK1 activation.

We next tested whether cells expressing methylation-deficient MCM4 experienced spontaneous DNA damage by quantifying chromatin-bound 53BP1 in G1 and S-phase cells (Figure 5D). The methylation-deficient MCM4 cells displayed a clear increase in chromatin-bound 53BP1 during G1 phase. One reasonable possibility is that single-strand DNA accumulation, as indicated by increased chromatin-bound RPA and RPA phosphorylation (Figure 5B and 5C), is the result of defective DNA replication which then results in under-replicated DNA accumulating in the subsequent G1 phase^69,70^.

To determine if DNA replication defects were evident in methylation-deficient *MCM4* mutants, we measured DNA replication fork rates using DNA combing (Figure 5E). Surprisingly, all three methylation-deficient MCM4 cell lines displayed increased fork rates, unlike cells depleted of PRMT1 where rates of DNA synthesis decreased (Figure 2C). Accelerated fork progression has been linked to reduced nascent-strand integrity and the accumulation of S1-sensitive daughter-strand gaps, prompting us to test whether the long tracts observed in methylation-deficient MCM4 cells similarly reflected gap-prone DNA replication^12,16,75,76^. We treated DNA from *MCM4* and *MCM4*_*Kall*_ cells with S1 nuclease prior to DNA combing (Figures 5F and S4C). S1 treatment markedly reduced the lengths of IdU labelled DNA tracts specifically from *MCM4*_*Kall*_ cells. Thus, we conclude that the accelerated fork rate observed in cells deficient in MCM4 arginine methylation reflects abnormal DNA replication with persistent ssDNA gaps.

Finally, we sought to define the relationship between the accelerated forks observed in the absence of PRMT1-dependent arginine methylation of MCM4 and the decreased fork rate observed when PRMT1 itself is depleted. We treated *MCM4* and *MCM4*_*Kall*_ cells with the PRMT1 inhibitor MS023 and measured DNA replication fork rates by DNA combing (Figures 5G and S4D). MS023 eliminated fork acceleration in *MCM4*_*Kall*_ cells, resulting in a decreased fork rate similar to that seen when wild-type cells were treated with MS023. We conclude that PRMT1 inhibition is epistatic to *MCM4*_*Kall*_ and infer that PRMT1 has replication-relevant targets in addition to MCM4.

### Defective arginine methylation alters the DNA replication fork-associated proteome

Given the pleiotropic nature of PRMT1 in regulating DNA replication, and that arginine methylation can alter protein localization^34^, we hypothesized that PRMT1 might regulate protein recruitment to replication forks. To measure changes in protein abundance at the replication fork we performed a comparative iPOND-MS experiment in *PRMT1* CRISPRi cells. We calculated the fold-change in spectral counts for each protein in the presence and absence of PRMT1 and ranked the results by z-score (Figures 6A and S5A). In agreement with our co-immunoprecipitation and AP-MS results (Figures S3E and 4G), MCM subunits were present at replication forks at similar levels independent of PRMT1 protein expression (Figure 6A, Table S1). Similarly, the recruitment of other core replisome components to replication forks was not substantially PRMT1-dependent (Table S1).

**Figure 6.**
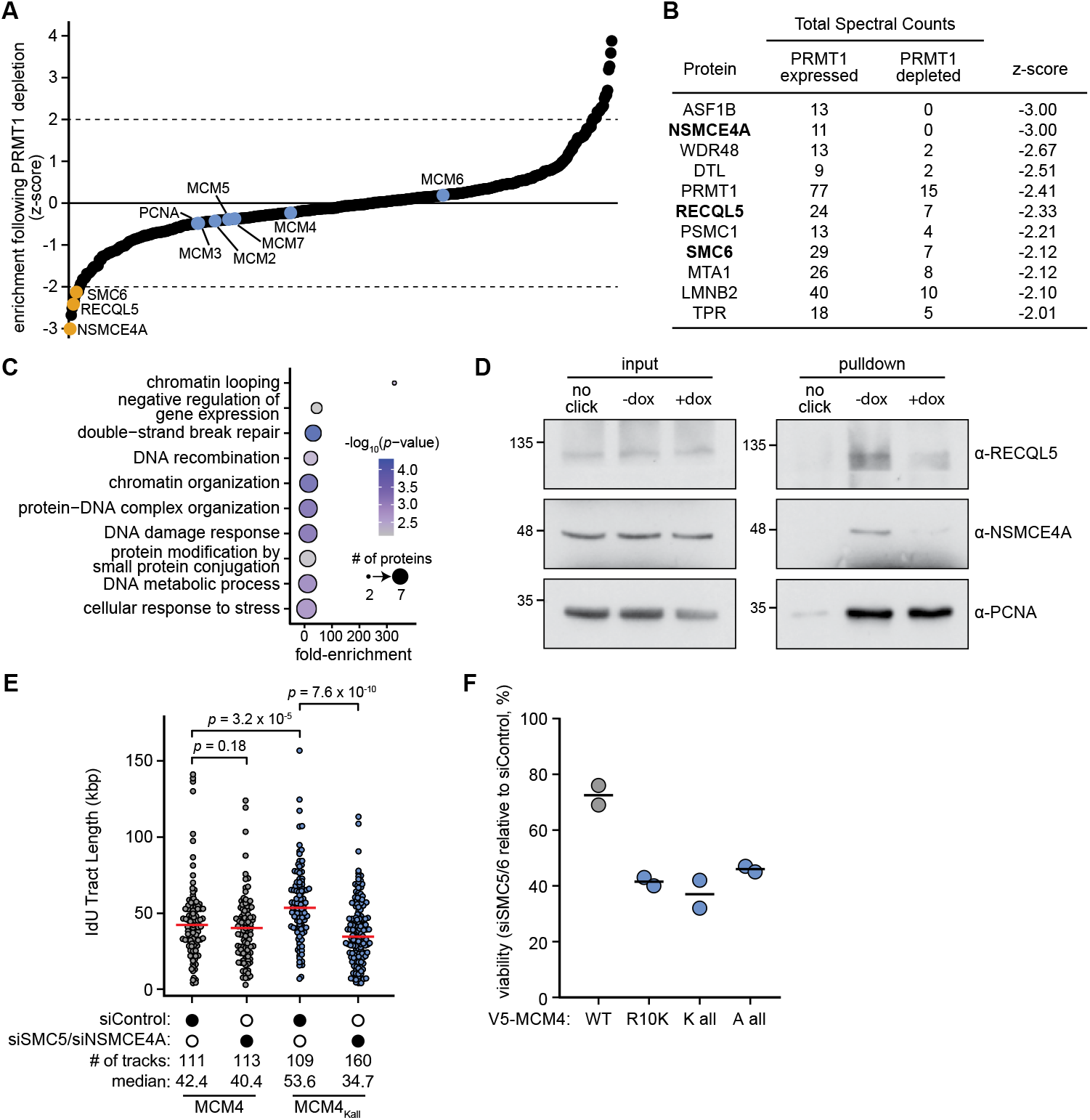
SMC5/6 is recruited to replication forks by PRMT1 and protects ssDNA gaps. (A) Quantification of replisome-associated proteins after *PRMT1* depletion. CRISPRi sgPRMT1-6 cells were treated with doxycycline for 5 days to deplete PRMT1, or mock treated, pulse labelled EdU for 15 minutes, fixed, and analyzed by iPOND-mass spectrometry. The fold-changes in protein spectral counts, in the EdU pulse, between PRMT1-depleted and untreated samples were averaged across three biological replicates and plotted with proteins ordered from lowest to highest z-score. Core replication proteins, MCM2-6 and PCNA, are highlighted in blue, and RECQL5, SMC6, and NSMCE4A are highlighted in orange. The dotted lines indicate z-scores of -2 and +2. *n*=3 biological replicates. (B) Table summarizing proteins in (A) with a z-score of ≤ -2. (C) Gene ontology (GO) biological process analysis for the 10 proteins in (B), excluding PRMT1. The fold-enrichment for each GO term is indicated, the colours indicate the corrected *p*-values, and the size of the circles corresponds to the number of genes annotated to the given GO term. (D) CRISPRi sgPRMT1-6 cells were treated with doxycycline for 5-days to deplete *PRMT1* (+dox), or mock treated (-dox), and subjected to the iPOND workflow. The input extracts and streptavidin pulldowns from the EdU pulse were immunoblotted for the presence of RECQL5, NSMCE4A, and PCNA. The control without covalent attachment of streptavidin to EdU (no click) is shown. The positions of molecular weight standards, in kDa, are indicated to the left. (E) DNA replication tract length analysis of *MCM4* knock-in lines following *SMC5/6* knockdown. *MCM4* and *MCM4*_*Kall*_ knock-in cells were treated with siRNAs targeting *NSMCE4A* and *SMC5*, followed by sequential pulse-labelling with CldU and IdU, DNA fibre isolation, and molecular combing. The IdU tract lengths were measured and are plotted. Medians are indicated by horizontal red bars and *p*-values were calculated with a two-sided Mann-Whitney *U* test. *n*=2 biological replicates (replicate 2, Figure S5C). (F) Viability of MCM4 knock-in lines following *SMC5/6* knockdown. *MCM4* and *MCM4*_*K all*_ knock-in cells were treated with siRNAs targeting NSMCE4A and SMC5. Cell viability was measured 4 days post-transfection and is plotted relative to cells transfected with control siRNA. *n*=2 biological replicates.

Contrasting with core replisome proteins, we identified 10 proteins that had decreased abundance at replication forks when PRMT1 was depleted (Figures 6A and 6B). Gene ontology analysis of the 10 proteins revealed strong enrichments for DNA repair and chromatin conformation functions (Figure 6C). RECQL5 and two members of the SMC5/6 complex, SMC6 and NSMCE4A, stood out as relevant to DNA replication fork integrity. RECQL5 displaces RAD51 during replication fork restart^77^. SMC5/6 can bind to ssDNA-dsDNA junctions^78^ and is enriched at interstrand crosslinks^79^, with both DNA structures being potential substrates for repriming. We confirmed the PRMT1 dependence of RECQL5 and NSMCE4A association with DNA replication forks using iPOND-western blotting (Figure 6D). Consistent with our mass spectrometry results (Table S1), PCNA abundance at replication forks was unchanged when PRMT1 was depleted, while the abundance of both RECQL5 and NSMCE4A were reduced. Importantly, the steady-state levels of RECQL5 and NSMCE4A were unchanged when PRMT1 was depleted (Figure 6D, input). Together, our data indicate that PRMT1 facilitates proper recruitment and/or maintenance of important fork stability proteins at replication forks.

### SMC5/6 protects ssDNA gaps following the loss of MCM4 arginine methylation

Recently, SMC5/6 was shown to protect replication-associated ssDNA gaps from breakage^80^. Since we find that members of SMC5/6 are less abundant at forks when PRMT1 is depleted, we hypothesized that ssDNA gaps present at replication forks in cells deficient in MCM4 arginine methylation could require SMC5/6 to prevent breakage. Indeed, siRNA knockdown of *SMC5* and *NSMCE4A* resulted in dramatically shorter DNA replication tracts in *MCM4*_*Kall*_ cells, suggesting that the SMC5/6 complex prevents breakage of the replication-associated ssDNA gaps that accumulate in the absence of MCM4 arginine methylation (Figures 6E, S5B, S5C, and S5D). Furthermore, SMC5/6 depletion in MCM4 methylation-deficient cells caused a synthetic fitness defect (Figure 6F). Thus, methylation-deficient MCM4 cells require SMC5/6 to maintain ssDNA gaps from aberrant DNA replication, and to sustain viability.

## Discussion

Despite major advances in defining the composition and function of the eukaryotic replisome, important regulators of DNA synthesis and fork integrity remain to be identified. Here we identify PRMT1 as a replisome-associated arginine methyltransferase that promotes both proper DNA synthesis and genome stability. PRMT1 localizes to active replication forks, its methyltransferase activity is required to sustain normal DNA synthesis, and its depletion or inhibition slows fork progression in multiple human cell types. We identify MCM4 as a direct PRMT1 substrate and show that loss of MCM4 arginine methylation causes aberrant DNA replication associated with persistent ssDNA gaps. In parallel, PRMT1 depletion alters the protein environment of the fork without broadly disrupting the core replisome, instead reducing the association of fork protection and repair factors including RECQL5 and the SMC5/6 complex. Together, our findings support a model in which PRMT1 preserves fork integrity through multiple coordinated mechanisms rather than through a single downstream effector (Figure 7).

**Figure 7.**
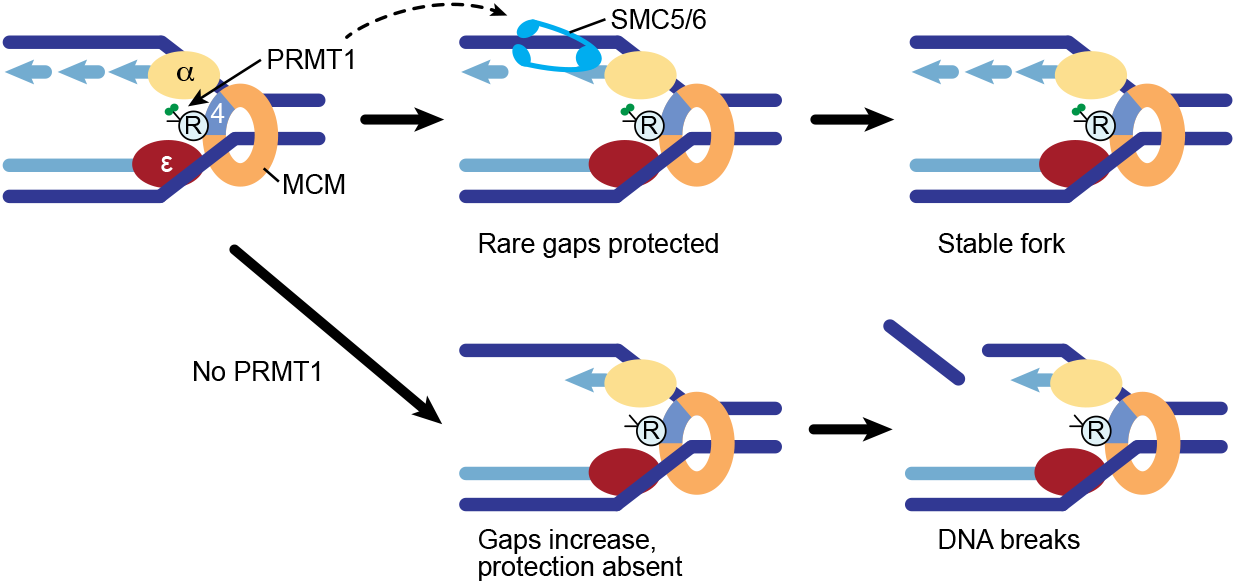
PRMT1 couples continuous DNA synthesis to stabilization of daughter-strand gaps. PRMT1 preserves fork integrity through two coordinated functions: methylation of MCM4 suppresses gap-prone DNA synthesis, and fork-associated SMC5/6 stabilizes vulnerable gap-containing intermediates. Loss of MCM4 methylation causes discontinuous DNA synthesis and daughter-strand gap accumulation, yielding apparent fork acceleration. Loss of PRMT1 combines this defect with reduced fork association of SMC5/6, destabilizing gap-containing intermediates and promoting breakage, replication stress, DNA damage, and loss of viability.

A central point emerging from our data is that the apparently divergent phenotypes of methylation-deficient MCM4 and PRMT1 depletion can be reconciled within one coherent model. Loss of MCM4 arginine methylation increases replication tract length, but this increase likely reflects only apparent fork acceleration. The accompanying increase in chromatin-bound RPA, activation of RPA32 Ser33 phosphorylation, and sensitivity of nascent DNA tracts to S1 nuclease indicate that DNA synthesis in the absence of MCM4 arginine methylation is gap-prone and discontinuous.

We infer that PRMT1-dependent methylation of MCM4 normally suppresses the formation of replication-associated ssDNA gaps, likely by coordinating productive fork progression and limiting aberrant repriming or uncoupling. Our inference is supported by prior work showing that the Mcm4 N-terminal regulatory domain integrates signaling inputs that influence initiation and fork progression^81–85^, and that the extreme amino terminus of human MCM4 contributes directly to MCM4/6/7 helicase activity^86^. Current structural work places the TIMELESS–TIPIN fork protection complex at the front of CMG, where TIMELESS engages the MCM N-tier, including MCM4^87^. Methylation of the MCM4 amino terminus could therefore modulate fork protection complex engagement or activity, with downstream consequences for helicase–polymerase coupling. In this framework, the longer tracts observed in methylation-deficient MCM4 cells would reflect faster apparent tract extension in the presence of ssDNA gaps, rather than faithful DNA replication.

Our data further indicate that MCM4 is not the only relevant target through which PRMT1 promotes DNA replication. Type I PRMT inhibition reduces fork rate in both wild-type and methylation-deficient MCM4 cells to similar levels, arguing that PRMT1 has additional functions beyond methylating MCM4. Consistent with that conclusion, comparative iPOND-MS shows that PRMT1 depletion leaves the abundance of core replisome components largely unchanged while selectively reducing the fork association of proteins linked to fork stability and repair. Among these, the loss of SMC5/6 is particularly notable, given established roles for SMC5/6 in replication fork stability and completion of DNA replication, particularly in difficult-to-replicate regions^88–90^, the preference of SMC5/6 for substrates resembling stressed forks^78,91^, and recent evidence that SMC5/6 protects replication-associated ssDNA gaps from breakage^80^. We show that SMC5/6 depletion in the context of methylation-deficient MCM4 shortens replication tracts and reduces viability. We propose an integrated model (Figure 7) in which PRMT1 promotes efficient fork-association or maintenance of SMC5/6, and SMC5/6 stabilizes the gap-containing fork intermediates that arise when MCM4 is not properly methylated. Thus, in the absence of PRMT1, two defects coincide: unmethylated MCM4 promotes formation of aberrant, gap-containing replication intermediates, while reduced fork association of SMC5/6 renders those intermediates vulnerable to breakage. The combined defect results in the short replication tracts we observe in PRMT1-deficient cells.

An important consequence of PRMT1 loss is accelerated genome instability. PRMT1-depleted cells display increased micronuclei, and elevated 53BP1-associated DNA damage during S/G2 and in G1 during aphidicolin treatment. We infer that PRMT1 deficiency generates replication-associated lesions during S-phase and allows a fraction of those lesions, or under-replicated regions^69,70^, to persist through mitosis into daughter cells. The methylation-deficient MCM4 mutants are informative in this regard: they show clear replication-stress phenotypes and increased G1-associated 53BP1 without a strong increase in S-phase 53BP1, consistent with ssDNA gaps persisting initially as incompletely replicated DNA and later becoming overt chromosome damage. Thus, PRMT1 protects the fidelity, not only the speed, of chromosome duplication.

Our results also suggest that PRMT1 contributes to the replication stress response at the level of checkpoint signaling, extending earlier work showing that PRMT1-dependent methylation of MRE11 supports the S-phase checkpoint and efficient ATR/CHK1 signaling^35,92^, as well as studies indicating that PRMT inhibition blunts replication-stress checkpoint responses in part by lowering ATR abundance^49,50^. Our data extend the emerging picture of PRMT1 function at stressed forks: ATR activation (increased RPA foci and RPA32 Ser 33 phosphorylation) is maintained when MCM4 is not PRMT1-methylated, but the checkpoint signal is not efficiently propagated to CHK1. PRMT1-deficient cells therefore experience a compound defect: they generate problematic replication intermediates while also mounting an incomplete checkpoint response to them. Although the mechanism connecting PRMT1 to CHK1 activation remains to be determined, it is of interest that MCM4 engages the fork protection complex (TIMELESS–TIPIN and CLASPIN), which is a mediator of efficient CHK1 activation at stalled forks ^93,94^. The selective checkpoint defect we observe likely contributes substantially to the spontaneous DNA damage and genome instability we observe upon PRMT1 loss.

In summary, our study expands the role of arginine methylation in genome maintenance beyond canonical DNA damage response factors to the control of active DNA replication forks. Previous work has shown that arginine methylation can regulate DNA repair enzymes and checkpoint proteins, but our data identify a direct replisome-centered role for PRMT1 in suppressing ssDNA gap formation and preserving the stability of vulnerable replication fork intermediates. Replication-associated ssDNA gaps are increasingly recognized as major sources of genome instability, mutagenesis, and therapeutic vulnerability^22,95^. Our findings therefore place PRMT1, MCM4 methylation, and SMC5/6-dependent daughter strand gap protection within a common pathway that limits endogenous replication stress. Although our results do not establish that this pathway fully explains the essentiality of PRMT1, they raise the possibility that one major function of PRMT1 in proliferating cells is to ensure that DNA synthesis is both continuous and structurally stable. More generally, our work supports a view of CMG as a regulatory platform whose activity can be shaped by arginine methylation at the MCM4 amino terminus, with consequences for fork architecture, checkpoint output, DNA synthesis, and the preservation of genome integrity.

## Materials and Methods

### Cell culture

HEK293T (ATCC), HEK293 (ATCC) and A375 (gift from Dr. Mikko Taipale) cells were maintained in Dulbecco’s modified Eagle’s medium (DMEM, Wisent) supplemented with 10% Fetal Bovine Serum (FBS, Wisent) and 1% penicillin/streptomycin (PS, Wisent). HAP1 cells were maintained in Iscove’s Modified Dulbecco’s Medium (IMDM) supplemented with 10% FBS and 1% PS. U2OS cells were maintained in McCoy’s 5A medium (Wisent) supplemented with 10% FBS and 1% PS.

To generate CRISPRi lines, A375 cells were first transduced with lentivirus to express dCas9-Zim3-mCherry^56^ and cell sorted on mCherry intensity. A375 cells with high mCherry signal were then infected with lentivirus to introduce the indicated doxycycline inducible sgRNA and selected with puromycin (3 µg/mL) for 2-4 days. All sgRNAs were induced by treating cells with doxycycline (100 ng/mL, refreshed every 2 days) for 5 days unless otherwise indicated.

For AP-MS of MCM4 variants, HEK293 Flp-In T-Rex cells were transfected with pOG44 vector and a pcDNA5 TO C-term UltraID vector containing the indicated *MCM4* variant and prepared as described previously^96^.

To generate knock-ins at the *MCM4* locus in HAP1 cells we used a modified qTAG protocol^97^. Cells were co-transfected with a px330-PITCh plasmid (gift from Dr. Laurence Pelletier, Addgene: 127875) containing a guide RNA targeting the 5’ end of the first exon of *MCM4* and a donor qTAG-N-Puro-V5 plasmid (gift from Dr. Laurence Pelletier, Addgene: 207697) consisting of two ∼900 bp homology arms flanking a V5-tag (see Key Resources Table). Four separate donor vectors containing mutations within the right homology arm to generate the appropriate *MCM4* allele were cloned (see Key Resources Table). Following transfection, cells were sub-cultured for 3 days and then treated with 1 µg/mL puromycin for 2 days. Monoclonal lines were generated by seeding single cells into 96-well plates. Clones homozygous for the appropriate knock-in were validated by PCR amplifying the 5’ end of *MCM4* from genomic DNA and Sanger sequencing (see Key Resources Table).

Cell lines were routinely monitored to ensure the absence of *Mycoplasma* contamination using the MycoAlert PLUS Mycoplasma Detection Kit (Lonza #LT07-705).

### Plasmid and siRNA transfections

For virus production, HEK293T cells were transfected with psPAX2 and pMD2.G packaging plasmids, and either a tet pLKO sgRNA puro vector or pHR UCOE SFFV dCas9 mCherry Zim3 KRAB using X-tremeGENE HP reagent (Roche). For all other plasmid transfections, cells were transfected using Lipofectamine 3000 Reagent. Lipofectamine RNAiMAX was used for siRNA transfections according to the manufacturer’s protocol.

### Constructs and cloning

To construct pcDNA3-N-FLAG vectors, *PRMT1* cDNA was PCR amplified. The pcDNA3-N-FLAG empty vector and PCR products were digested with EcoRI and BamHI, and ligated. The *PRMT1*^*E153Q*^ pcDNA3-N-FLAG vector was constructed by QuickChange site-directed mutagenesis.

pDONR223 vectors containing *MCM2, MCM3* or *MCM5* were acquired from the human ORFeome v8.1 library. *MRE11* (pcDNA5 MRE11), *MCM4* (pACT2 MCM4) and *MCM6* (pACT2 MCM6) cDNAs were PCR amplified and assembled into the pDONR221 vector using a Gateway BP reaction. pENTR221 MCM7 was assembled into the pDONR221 vector using a Gateway BP reaction. Donor vectors were then assembled into either pcDNA-DEST40 or pcDNA3.1-ccdB-eGFP destination vectors using a Gateway LR reaction.

To construct pGEX-6P-1 bacterial expression vectors, cDNA for *MCM4, PRMT1, PRMT1*_*E153Q*_ and residues 566-600 of *MRE11* were PCR amplified. The pGEX-6P-1 vector and PCR products were digested with BamHI and XhoI, and ligated.

The pDONR221 *MCM4*_*R10K*_ vector was constructed by QuickChange site-directed mutagenesis. pDONR221 *MCM4*_*R14K*_ was constructed by PCR amplifying *MCM4* cDNA with mutagenic primers. A second PCR amplification was performed to incorporate attB sites onto the *MCM4*_*R14K*_ amplicon before assembly into pDONR221 using a Gateway BP reaction. To construct pDONR221 *MCM4*_*Kall*_ and pDONR221 *MCM4*_*Aall*_, *MCM4* cDNA was PCR amplified with mutagenic primers. The pDONR221 *MCM4*_*WT*_ vector and PCR products were digested with SalI and NotI, and ligated. pDONR221 vectors were assembled into pcDNA-DEST40 or pcDNA5 TO C-term UltraID destination vectors using a Gateway LR reaction. The pcDNA3.1 nV5-DEST MCM4 plasmids were constructed by PCR amplifying *MCM4* cDNA. The pcDNA3.1 nV5-DEST vector was digested with EcoRI and BsrGI, and PCR products were digested with EcoRI and BsaI, and ligated.

sgRNAs were cloned into the tet pLKO sgRNA puro and px330-BbsI-PITCh^97^ vectors as described previously. Repair template qTAG-N-Puro-V5 vectors were cloned by Gibson assembly as described previously^97^. The left homology arm was generated by PCR amplifying HAP1 genomic DNA. Right homology arms, summarized in the Key Resources Table, were synthesized and PCR amplified.

### EdU secondary screen

U2OS cells were transfected in clear-bottom 96-well plates (Fisher Scientific) with 40 nM siRNA for 48 hours, using Lipofectamine RNAiMAX (Life Technologies). A negative non-targeting control siRNA (Dharmacon, GE Healthcare) and a siGENOME SMARTpool mixture of four siRNAs targeting each gene (Dharmacon, GE Healthcare) were used in the screen. 10 μM EdU was added to the culture media for the last 30 mins post-transfection. Cells were washed with PBS, fixed with 4% paraformaldehyde for 10 min and permeabilized with 0.3% Triton X-100 for 10 minutes. EdU was detected by conjugation to Alexa-Fluor 488 azide (Life Technologies). Cells were blocked with PBS buffer containing 10% goat serum, 0.5% NP-40 and 0.5% saponin for 30 minutes at room temperature. After washing with PBS, cells were incubated in DAPI (0.5 µg/mL) for 1 hour at room temperature. Images were acquired from each well on an IN Cell Analyzer 6000 (GE Healthcare Life Sciences) at 10x magnification at non-saturating exposure times. Columbus imaging software (PerkinElmer) was used to segment nuclei with the DAPI channel and the mean EdU signal intensity per nucleus was calculated.

### EdU flow cytometry

For EdU flow cytometry experiments, cells were pulse labelled with 10 µM (A375) or 20 µM (U2OS) EdU for 30 minutes then fixed in 4% paraformaldehyde for 15 minutes at room temperature. EdU was then fluorescently labelled as described in the Click-iT EdU Alexa Fluor 647 Flow Cytometry Assay Kit (Thermo). Samples were then incubated in 0.5 µg/mL 4′,6-diamidino-2-phenylindole (DAPI) and 250 µg/mL RNase A for 30 minutes at room temperature prior to being analyzed on either a BD LSRII or BD LSR Fortessa cytometer using FACS Diva software (BD Biosciences). For the PRMT1 rescue experiment, A375 cells were transfected with a pcDNA3 N-FLAG vector carrying *PRMT1* or *PRMT1*_*E153Q*_. Flow cytometry samples were prepared 48 hours post-transfection as described above.

To quantify chromatin bound 53BP1 or RPA32, cells were pulse labelled with 10 µM (HAP1) or 20 µM (U2OS) EdU for 30 minutes then incubated in ice cold cytoskeletal (CSK) buffer (300 mM sucrose, 100 mM NaCl, 3 mM MgCl_2_, 10 mM PIPES pH 7.0, 0.5% Triton X-100) containing cOmplete Protease Inhibitor Cocktail (Roche) for 5 minutes at 4°C. Cells were then pelleted, washed in 1% (wt/vol) BSA in PBS (PBS-B) and fixed in 4% paraformaldehyde for 15 minutes at room temperature. Fixed samples were incubated in blocking buffer (0.1% NP-40 in PBS-B) for 10 minutes before incubation in primary antibody in blocking buffer, overnight at 4°C. Samples were then washed with blocking buffer and incubated with secondary antibody for 30 minutes at room temperature. EdU was conjugated to Alexa Fluor 647 as described in the Click-iT EdU Alexa Fluor 647 Flow Cytometry Assay Kit (Thermo), and samples were washed and incubated in 0.5 µg/mL DAPI and 250 µg/mL RNase A for 30 minutes at room temperature prior to being analyzed on either a BD LSRII or BD LSR Fortessa cytometer using FACS Diva software (BD Biosciences).

### DNA combing

Prior to DNA combing, cells were treated with doxycycline for 5 days (CRISPRi PRMT1 depletion), with 0.1, 0.25, 0.5, or 2 µM MS023 for 48 hours (PRMT1 inhibition), or left untreated. Exponentially growing cells were labelled with 25 µM CldU in culture medium for 30 minutes, washed briefly with PBS and labelled with 125 µM IdU in culture medium for 30 minutes. Cells were embedded in low melting point agarose (0.5% agarose final) and digested in Proteinase K buffer (1 mg/mL Proteinase K, 500 mM EDTA, 1% sarkosyl) for three days at 50°C protected from light. Proteinase K buffer was refreshed every 24 hours. Agarose plugs were melted and DNA fibres were stretched on silanized coverslips and labelled by immunofluorescence as described previously^58^ with the following modifications. After blocking, the coverslips were incubated in blocking buffer containing primary antibodies (anti-IdU, and anti-CldU) for 1 hour at 37°C. Coverslips were washed three times in PBS-T (0.05% vol/vol) and incubated in blocking buffer containing anti-DNA antibody for 1 hour at 37°C. Coverslips were washed three times in PBS-T and incubated in blocking buffer containing secondary antibody (anti-rat Alexa Fluor 488, anti-mouse IgG1 Alexa Fluor 546, anti-mouse IgG2a Alexa Fluor 647). DNA fibres were imaged on an Imager. Z1 fluorescence microscope (Zeiss) with a 63x objective and IdU tract lengths were measured using Fiji^98^ and are expressed as kbp or kbp/min.

The S1 nuclease DNA combing assay was performed as described previously^13^, except that plugs were treated with 15U of S1 nuclease for 1 hour.

### Quantification of micronuclei

A375 cells were seeded onto coverslips and cultured for 5 days in the presence or absence of doxycycline to deplete PRMT1. Cells were then briefly washed in ice cold PBS and nuclear membranes were permeabilized by treating samples with ice cold nuclear extraction buffer (20 mM HEPES pH 7.5, 20 mM NaCl, 5 mM MgCl_2_, 1 mM DTT, 0.5% NP-40, cOmplete Protease Inhibitor Cocktail) for 10 minutes on ice. Cells were fixed in 2% formaldehyde for 10 minutes at room temperature, washed with PBS, and treated with 0.4 µg/mL DAPI. Prior to imaging, cover slips were washed three times with PBS and mounted on microscope slides with Prolong Gold.

To image micronuclei, a minimum of 15 fields per sample were randomly imaged on a Nikon Eclipse Ti2 inverted microscope. The number of nuclei per sample was quantified using CellProfiler and micronuclei were counted manually. Micronuclei incidence was reported as a percentage and calculated by dividing the number micronuclei per sample by the total number of nuclei. A minimum of 730 nuclei (cells) were counted per condition per replicate.

### 53BP1 immunofluorescence

To quantify 53BP1 foci, A375 CRISPRi cells were seeded into black 96-well plates and grown for 5 days in the presence or absence of 100 ng/mL doxycycline. To induce replication stress, cells were treated with 0.4 µM aphidicolin for 16 hours prior to fixation. Cells were then fixed in 4% paraformaldehyde in PBS for 10 minutes at room temperature before being permeabilized with 0.5% Triton X-100 in PBS for 10 minutes. Samples were then incubated in blocking buffer (5% BSA and 0.05% Tween-20 in PBS) for 1 hour. Primary and secondary antibodies were diluted in blocking buffer and added to cells. Cells were first incubated with anti-cyclin A antibody, then anti-mouse IgG2A Alexa Fluor 647 antibody, then mouse anti-53BP1 antibody, and finally anti-mouse IgG1 Alexa Fluor546 antibody. All antibody incubations lasted 1 hour, except for the 53BP1 antibody which was incubated for 1.5 hours. Fixed samples were washed three times with PBS between each antibody addition. Lastly, cells were treated with 0.15 µg/mL DAPI for 20 minutes, washed three times with PBS and imaged on an Opera Phenix confocal microscope. Images were analyzed using CellProfiler. Cells negative for cyclin A staining were classified as being in G1 and cells positive for cyclin A staining were classified as being in S-phase or G2. A minimum of 3,000 cells were analyzed per condition for each replicate.

### RPA immunofluorescence

To quantify RPA foci, A375 CRISPRi cells were seeded into black 96-well plates and grown for 5 days in the presence or absence of 100 ng/mL doxycycline. To label S-phase cells, 10 µM EdU was added to the culture medium for 20 minutes prior to the addition of 0, 1.9, 7.5, or 30 µM aphidicolin for 30 minutes. Cells were then washed briefly with ice cold PBS and CSK buffer without Triton X-100. To remove proteins which are not chromatin bound, samples were then incubated in CSK buffer on ice for 15 minutes, washed with PBS, and then fixed in 4% paraformaldehyde in PBS for 15 minutes at room temperature. EdU was fluorescently labelled as described in the Click-iT EdU Alexa Fluor 647 Flow Cytometry Assay Kit (Thermo). Samples were then briefly washed with PBS and incubated in blocking buffer (10% goat serum, 0.5% NP-40 and 0.5% saponin in PBS) for 1 hour at room temperature. Subsequently, cells were incubated with an anti-RPA32 primary antibody for 2 hours at room temperature, washed with PBS, then incubated with anti-mouse Alexa Fluor 488 secondary antibody for 1 hour at room temperature. Lastly, cells were treated with 0.15 µg/mL DAPI for 20 minutes, washed three times with PBS and imaged on an Opera Phenix confocal microscope. Images were analyzed using CellProfiler. Only cells positive for EdU staining were considered in the analysis. A minimum of 88 S-phase cells were analyzed per condition for each replicate.

### Whole cell extracts

Whole cell extracts were prepared by incubating cells in RIPA buffer (10 mM Tris pH 8.0, 150 mM NaCl, 1% Triton X-100, 0.1% sodium deoxycholate, 0.1% SDS, 1 mM EDTA, 1 mM DTT) containing cOmplete Protease Inhibitor Cocktail and phosSTOP (Roche) for 15 minutes on ice prior to centrifugation at 16,100 x *g* for 15 minutes at 4°C. Supernatants were then isolated and protein concentration was normalized by Bradford assay (Sigma).

### Nuclear fractionation

To extract nuclear proteins, cells were first resuspended in hypotonic lysis buffer (50 mM Tris pH 7.5, 10 mM NaCl, 1.5 mM MgCl_2_, 10% glycerol, 0.34 M sucrose, 1 mM DTT) containing cOmplete Protease Inhibitor Cocktail and phosSTOP for 10 minutes with gentle shaking at 4°C. Samples were then incubated for an additional 10 minutes on ice prior to the addition of Triton X-100 to a final concentration of 0.1% (vol/vol) and incubated for another 10 minutes. Nuclei were then sedimented by centrifugation at 1,300 g for 5 minutes at 4°C. The supernatant (cytoplasmic fraction) was removed, and cell nuclei were washed once with hypotonic lysis buffer, sedimented, and resuspended in nuclear extraction buffer (50 mM Tris pH 7.5, 420 mM NaCl, 10% glycerol, 1 mM EDTA, 1 mM DTT) containing cOmplete Protease Inhibitor Cocktail and phosSTOP and incubated for 1 hour at 4°C with agitation. To ensure complete nuclear lysis, samples were sonicated for 10 seconds (Branson Model 250, tapered microtip; constant duty cycle and output setting 2) then centrifuged at 16,100 x *g* for 15 minutes at 4°C. The soluble fraction (nuclear fraction) was then isolated, and protein concentration was measured by Bradford assay (Sigma).

### Immunoprecipitations

All immunoprecipitations were performed using nuclear extracts prepared as described above. Magnetic Protein G Dynabeads beads were washed twice in nuclear extraction buffer then conjugated to the appropriate antibody (1 µg anti-V5 antibody/IP, 3 µL anti-GFP antibody/500 µL nuclear extract) by mixing and incubating overnight at 4°C with agitation. Protein A Dynabeads were mixed with anti-MCM2 antibody (5 µg antibody/IP) and incubated overnight at 4°C with agitation. The next day, antibody bound Dynabeads were washed twice with nuclear extraction buffer prior to being resuspended in the appropriate nuclear extract and incubated overnight at 4°C with agitation. Pulldowns were then washed an additional two times in nuclear extraction buffer before proteins were eluted by resuspending the Dynabeads in 2x Laemmli sample buffer and boiling at 95°C for 10 minutes. The samples were then briefly centrifuged, and supernatants were analyzed by immunoblotting.

### Immunoblot analysis

Cell extracts were added to 4X Laemmli sample buffer and boiled at 95°C for 10 minutes. Proteins were resolved by SDS-PAGE, transferred to nitrocellulose membranes, blocked for 1 hour in 5% (w/v) milk powder in Tris-buffered saline with 0.5% Tween-20 (TBS-T), and probed with primary antibodies overnight at 4°C. Membranes were then washed three times in TBS-T, probed with secondary antibodies for 1 hour at room temperature and washed an additional three times in TBS-T. Protein expression was detected using SuperSignal West Pico PLUS chemiluminescent substrate (Thermo Fisher) and visualized on X-ray film or a ChemiDoc (BioRad). Blots using the rabbit anti-pCHK1 (Ser 345) antibody were performed as described above except 5% (w/v) BSA replaced milk in the blocking buffer.

### Protein purification

BL21(DE3)pLysS was transformed with pGEX vectors encoding GST-tagged PRMT1, PRMT1_E153Q_, GAR, or MCM4. Cultures were induced with 0.2 mM IPTG and incubated overnight at 30°C with vigorous shaking. Bacteria were pelleted by centrifugation, resuspended in bacterial resuspension buffer (25 mM Tris pH 7.6, 300 mM NaCl, 10% Glycerol, 0.2 mM EDTA, 5 mM β-mercaptoethanol, 0.5% NP-40) supplemented with 1 mM PMSF, and flash frozen in liquid nitrogen. Samples were thawed at 37°C and the freeze thaw cycle was repeated two more times. Lysates were sonicated in 15 seconds bursts followed by 10 seconds rest using a Misonix S4000 at 20% amplitude over a minute and 45 seconds on ice. Insoluble material was removed by centrifugation at 30,000 x *g* for 30 minutes at 4°C.

The GST tagged proteins were purified using a GSTrap FF column (GE Healthcare) and Äkta Pure system (GE Healthcare). Lysates were injected onto the system, the column was washed with 3 column volumes of bacterial resuspension buffer, and bound proteins were eluted in elution buffer (20 mM Tris pH 7.6, 100 mM NaCl, 0.2 mM EDTA, 20% glycerol, 5 mM DTT, 10 mM glutathione). Samples were dialyzed thrice against BC100 (10 mM Tris pH 7.6, 100 mM NaCl, 10% glycerol, 5 mM EDTA, 1 mM DTT) and analyzed by SDS-PAGE followed by Coomassie blue staining. MCM4 was eluted through on-column cleavage of the GST tag using PreScission protease (Genscript) at 2U/100 µg of GST-MCM4 protein.

### Methylation assays in vitro

Methylation assays were performed as described previously^99^. 150 nM recombinant GST-PRMT1 (wild type or E153Q) was incubated with 3 µM recombinant MCM4 in PBS containing 500 µM S-adenosylmethionine (NEB) for 3 hours at 30°C. Reactions were halted by adding 4x Laemmli sample buffer to a final concentration of 1x and boiling for 10 minutes at 95°C. Arginine methylation was detected by immunoblotting.

### Cell viability assays

Cells were transfected with the indicated siRNA and then seeded into 96-well plates at equal density. 4 days post-transfection, cell viability was analyzed with CellTiter-Glo (Promega), and luminescence was measured on a CLARIOstar Plus microplate reader.

### iPOND sample preparation

iPOND experiments were performed essentially as described previously^52^. To label nascent DNA, cells were pulsed with either 10 µM EdU for 10 minutes (HEK293) or 12.5 µM EdU for 15 minutes (A375). When a thymidine chase was applied, cells were briefly washed with PBS containing 10 µM thymidine then incubated in culture medium containing 10 µM thymidine for 30 minutes (HEK293) or 12.5 µM thymidine for 1 hour (A375). Either 1.5 × 10^8^ (HEK293) or 2 × 10^8^ (A375) cells from each sample were cross-linked with 1% formaldehyde (Sigma, 252549) in PBS for 20 minutes and quenched with 0.125 M glycine. Cells were washed three times in PBS and permeabilized with 0.25% Triton X-100 in PBS for 30 minutes. Cell pellets were then washed once with 0.5% BSA in PBS and once with PBS prior to the conjugation of EdU with biotin azide (HEK293: 1 µM, Life Technologies; A375: 10 µM, Thermo) using Click-iT reaction mix (HEK293: Invitrogen) or 2 mM copper sulfate and 100 mM sodium ascorbate in PBS (A375) for 2 hours. Cells were washed once with 0.5% BSA in PBS and then resuspended in lysis buffer (1% SDS, 50 mM Tris pH 8.0) containing cOmplete Protease Inhibitor Cocktail. Samples were sonicated using five cycles of a 20 second pulse and a 40 second pause, on ice (Branson Model 250, tapered microtip, constant duty cycle, output setting 2). Lysates were centrifuged at 14,000 rpm for 20 minutes at 4°C and supernatants were diluted 1:1 (vol/vol) in PBS containing cOmplete Protease Inhibitor Cocktail. Affinity purification was performed by mixing lysates with streptavidin-agarose beads (HEK293: Novagen; A375: Millipore) and incubating the mixtures overnight at 4°C with rocking. When performing immunoblotting, beads were washed twice with lysis buffer, once with 1 M NaCl, and once more with lysis buffer before being resuspended in 2x Laemmli sample buffer and incubated for 30 minutes at 95°C.

### HEK293 iPOND mass spectrometry

To analyze iPOND streptavidin pulldowns by mass spectrometry, lysates were centrifuged at 16,100 x *g* or 20 minutes at 4°C and 900 µl of supernatants was diluted in 2 mL RIPA buffer (50 mM Tris-HCl pH 7.5, 150 mM NaCl, 1% Triton X-100, 1 mM EDTA, 1 mM EGTA, 0.1% SDS, 0.5% sodium deoxycholate) containing cOmplete Protease Inhibitor Cocktail. Pulldowns were performed by mixing lysates with 30 µl bed volume of streptavidin-sepharose beads (Cytiva, pre-washed 3 times with 1mL RIPA buffer) and incubating the mixtures overnight at 4°C with agitation. The beads pelleted (400 x *g*, 1 min), the supernatant removed, and the beads transferred to a 1.5 mL microfuge tube in 1 mL RIPA buffer (minus protease inhibitors and sodium deoxycholate). The beads were then washed by pipetting up and down (4x per wash step) first with an additional 1 mL RIPA buffer (minus protease inhibitors and sodium deoxycholate) followed by two washes in TAP lysis buffer (50 mM HEPES-KOH pH 8.0, 100 mM KCl, 10% glycerol, 2 mM EDTA, 0.1% NP-40), then 3 washes in 50 mM ammonium bicarbonate pH 8 (ABC). Beads were pelleted by centrifugation (400 x g, 1 min) and the supernatant aspirated in between wash steps. After the last wash, all residual 50 mM ABC was pipetted off and beads were re-suspended in 200 µL of 50 mM ammonium bicarbonate (ABC) pH 8 with 1 µg trypsin added and incubated at 37°C overnight with mixing. The next day an additional 0.5 µg of trypsin was added to each sample (in 10 µL 50 mM ABC) and the samples incubated an additional 2 hours at 37°C with mixing. Beads were then pelleted (400 x g, 2 min) and the supernatant transferred to a fresh 1.5 mL microfuge tube. The beads were then rinsed 2 times with 30 µL of HPLC H_2_O each time (pelleting beads at 400 x g, 2 min in between) and these rinses combined with the original supernatant. The pooled supernatant was then centrifuged at 16,100 x g for 10 minutes and most of the supernatant (minus 30 µL to exclude all beads) was transferred to a new 1.5 mL tube. Samples were then vacuum dried. For mass spectrometry acquisition, samples were resuspended in 12 µL of 5% formic acid and 5 µL was injected.

Nano-spray emitters were generated from fused silica capillary tubing, with 75 µm internal diameter, 365 µm outer diameter and 5-8 µm tip opening, using a laser puller (Sutter Instrument Co., model P-2000, with parameters set as heat: 280, FIL = 0, VEL = 18, DEL = 200). Nano-spray emitters were packed with C18 reversed-phase material (Reprosil-Pur 120 C18-AQ, 3 µm) resuspended in methanol using a pressure injection cell. 5 µL of sample in 5% formic acid was directly loaded at 800 nL/min for 20 min onto a 100 µm x15 cm nano-spray emitter. Peptides were eluted from the column with an acetonitrile gradient generated by a NanoLC-Ultra 2D plus HPLC system (Eksigent, Dublin, USA) and analyzed on an LTQ-Orbitrap Velos or the Orbitrap Elite (Thermo Electron, Bremen, Germany) equipped with a nanoelectrospray ion source (Proxeon Biosystems, Odense, Denmark). The LTQ-Orbitrap Velos and Orbitrap Elite instrument under Xcalibur 2.0 was operated in the data dependent mode to automatically switch between MS and up to 10 subsequent MS/MS acquisition. Buffer A is 100% H_2_O, 0.1% formic acid; buffer B is 100 ACN, 0.1% formic acid. The HPLC gradient program delivered an acetonitrile gradient over 125 minutes. For the first twenty minutes, the flow rate was 400 µL/min at 2% B. The flow rate was then reduced to 200 µL/min and the fraction of solvent B increase in a linear fashion to 35% until the 95.5 minutes. Solvent B was then increased to 80% over 5 minutes and maintained at that level until 107 minutes. The mobile phase was then reduced 2% B until the end of the run (125 min). The parameters for data dependent acquisition on the mass spectrometer were: 1 centroid MS (mass range 400-2000) followed by MS/MS on the 10 most abundant ions. General parameters were: activation type = CID, isolation width = 1 m/z, normalized collision energy = 35, activation Q = 0.25, activation time = 10 msec. For data dependent acquisition, minimum threshold was 500, the repeat count = 1, repeat duration = 30 sec, exclusion size list = 500, exclusion duration = 30 sec, exclusion mass width (by mass) = low 0.03, high 0.03.

MS data generated were stored, searched, and analyzed using the ProHits laboratory information management system (LIMS) platform and searched using Mascot (v2.3.02) and Comet (v2016.01 rev.2). The spectra were searched with the human and adenovirus sequences in the RefSeq database (version 57, January 30th, 2013) acquired from NCBI, supplemented with “common contaminants” from the Max Planck Institute (http://www.coxdocs.org/doku.php?id=maxquant:start_downloads.htm) and the Global Proteome Machine (GPM; ftp://ftp.thegpm.org/fasta/cRAP), and forward & reverse sequences (labeled “gi|9999” or “DECOY. Database parameters were set to search for tryptic cleavages, allowing up to 2 missed cleavages sites per peptide with a mass tolerance of 12 ppm for precursors with charges of 1+ to 3+ and a tolerance of 0.6 Da for fragment ions. Results from each search engine were analyzed through TPP (the Trans-Proteomic Pipeline, v.4.7 POLAR VORTEX rev 1) via the iProphet pipeline. Files were analyzed with SAINTexpress v3.1., the no CLK conditions was used as the control for DDA SAINT analyses. In both, replicates for controls and baits were set to 2.

### HEK293 iPOND-MS data analysis

Raw spectral count data was analyzed by the Significance Analysis of INTeractome (SAINT) label free quantification (LFQ) computational analysis tool^100^. Prey proteins with a SAINT score ≥ 0.95, at least 6 spectral counts across two biological replicates in the EdU pulsed sample, and a ≥ 2-fold enrichment in spectral counts in the EdU pulse relative to the thymidine chase were considered positive iPOND candidates. Positive iPOND candidates were plotted as a STRING network^54^ (accessed April 9^th^, 2026) with nodes representing proteins and edges marking high-confidence protein-protein interactions.

### PRMT1 comparative iPOND mass spectrometry

Samples were processed for streptavidin affinity purification and on-bead tryptic digestion as described above for HEK293 iPOND mass spectrometry. For data-dependent acquisition LC-MS/MS, one-quarter of each peptide sample was analyzed by nano-HPLC coupled to mass spectrometry. Nano-spray emitters were prepared and packed with C18 reversed-phase material as described above, except that fused silica capillaries with a 100 µm internal diameter and 365 µm outer diameter were used, and the laser puller delay parameter was set to DEL = 2000. Peptides resuspended in 5% formic acid were directly loaded at 800 nL/min for 20 min onto a 100 µm × 15 cm nano-spray emitter.

Peptides were eluted using an acetonitrile gradient generated by an Eksigent ekspert− nanoLC 425 and analyzed on a TripleTOF− 6600 instrument (AB SCIEX, Concord, Ontario, Canada). The gradient was delivered at 400 nL/min from 2% acetonitrile/0.1% formic acid to 35% acetonitrile/0.1% formic acid over 90 min, followed by a 15-min wash at 80% acetonitrile/0.1% formic acid and a 15-min equilibration to 2% acetonitrile/0.1% formic acid. The total DDA protocol was 135 min. MS1 scans were acquired with an accumulation time of 250 ms over a mass range of 400–1800 Da, followed by MS/MS scans of the top 10 candidate ions with an accumulation time of 100 ms per MS/MS scan. Candidate ions were required to have charge states of 2+ to 5+ and a minimum intensity threshold of 300 counts/s, were isolated using a 50 mDa window, and were dynamically excluded for 7 s after prior analysis.

Mass spectrometry data were stored, searched, and analyzed using the ProHits LIMS platform as described above. For this dataset, WIFF files were converted to MGF format using WIFF2MGF and to mzML format using ProteoWizard v3.0.10702 and the AB SCIEX MS Data Converter v1.3 beta. Searches were performed using Mascot v2.3.02 and Comet v2016.01 rev.2 against the human and adenovirus RefSeq database, supplemented with common contaminant databases, forward and reverse decoy sequences, sequence tags for BirA, GST26, mCherry, and GFP, and streptavidin, for a total of 72,481 entries. Search parameters allowed tryptic peptides with up to two missed cleavages, precursor mass tolerance of 35 ppm for charges 2+ to 4+, and fragment ion tolerance of 0.15 amu. Variable modifications included deamidation of asparagine and glutamine and oxidation of methionine. Search results were analyzed through the Trans-Proteomic Pipeline v4.7 POLAR VORTEX rev 1 using the iProphet pipeline.

### PRMT1 comparative iPOND-MS data analysis

Raw spectral count data was first analyzed by filtering for proteins with at least 1spectral count in the -dox EdU pulse (PRMT1 expressed) sample for each biological replicate. Frequent false-positive proteins were removed by filtering the hit list for proteins identified in >60% of standardized negative control AP-MS experiments in the CRAPome^101^ (accessed September 23^rd^, 2023). The fold-change of spectral counts for each biological replicate was calculated for each protein and the average fold-change for the +dox EdU pulse (PRMT1 depleted) was divided by the average fold-change for the -dox EdU pulse (PRMT1 expressed) samples. The fold-changes were used to calculate a z-score, and a one-sided paired Student’s t-test was applied to proteins depleted at the replication fork following PRMT1 depletion (z <-2) and proteins enriched at the replication fork following PRMT1 depletion (z > -2).

### Affinity purification-mass spectrometry of MCM4 variants

HEK293 Flp-In T-Rex cells were transfected with pOG44 vector and a pcDNA5 TO C-term UltraID vector containing the indicated *MCM4* variant. 3 days post-transfection, successful integrants were selected by treating cells with 200 µg/mL hygromycin B for 10 days. Cells were then seeded into 15-cm dishes and upon reaching 90% confluency, MCM4 variant expression was induced by treating cells with 1 µg/mL doxycycline for ∼16 hours. Plates were washed once with PBS, and cell pellets were collected in 1 mL PBS using a cell scraper. Samples were normalized by pellet weight and lysed as described previously^96^. FLAG-pulldowns, preparation of samples for mass spectrometric analysis and the subsequent analysis was performed as described previously^96^.

## Supporting information

Key Resources Table

Numerical Source Data

## Data availability

Data has been deposited as a complete submission to the MassIVE repository (https://massive.ucsd.edu/ProteoSAFe/static/massive.jsp) and assigned the accession number MSV000100773.

## Acknowledgements

The authors thank Stéphane Richard for helpful discussions and for providing ASYM26 antibody, members of the Brown Lab for valuable comments on the manuscript, Laurence Pelletier for providing px330-PITCh and qTAG-N-Puro-V5 plasmids, Mikko Taipale for providing pcDNA3.1 nV5 DEST, pcDNA 3.1 ccdB EGFP and pHR UCOE SFFV dCas9 mCherry Zim3 KRAB plasmids, Haley Wyatt for providing pGEX-6P1 plasmid, Lori Frappier for providing pACT2 MCM4 and pACT2 MCM6 plasmids, and Dan Durocher for providing the pcDNA5 MRE11 plasmid. MWF was supported in part by an OGS award from the Government of Ontario and a PGSD award from the Natural Sciences and Engineering Research Council of Canada. AM was supported by an Ontario Trillium Scholarship. SMM was supported by a University of Toronto Open Scholarship. EIC was supported by the Canadian Institutes of Health Research (RN508231–497056) and the Natural Sciences and Engineering Research Council of Canada (RGPIN-2024-06709). ACG is a Tier 1 Canada Research Chair in Functional Proteomics. Proteomics work was performed at the Network Biology Collaborative Centre (RRID: SCR_025375) at the Lunenfeld-Tanenbaum Research Institute, a facility supported by Canada Foundation for Innovation and the Government of Ontario. GWB is a Tier I Canada Research Chair in Genome Integrity and was supported by the Canadian Institutes of Health Research (FDN-159913 and RN566824-540094). We are grateful to work on the lands of the Mississaugas of the Credit, the Anishnaabeg, the Haudenosaunee, and the Wendat peoples, land that is now home to many diverse First Nations, Inuit, and Métis peoples.

## Author Contributions

MWF: Conceptualization, Formal analysis, Investigation, Methodology, Formal analysis, Writing-original draft, Writing-review & editing. CJW: Investigation, Formal analysis, Writing-review & editing. XY: Investigation. EC: Investigation, Writing-review & editing. AM: Investigation. SMM: Investigation. WHD: Investigation. Z-YL: Investigation. EIC: Formal analysis, Funding acquisition, Supervision, Writing-review & editing. ACG: Formal analysis, Funding acquisition, Supervision, Writing-review & editing. GWB: Conceptualization, Formal analysis, Funding acquisition, Supervision, Writing-original draft, Writing-review & editing.

