## Supplemental Table and Figures for "PRMT1 arginine methylation of MCM4 restricts ssDNA gap formation during DNA replication"

**Table S1. Replication protein abundance at replication forks in the presence and absence of PRMT1**

| protein | total spectral counts ( EdU pulse) PRMT1 expressed | total spectral counts ( EdU pulse) PRMT1 depleted | z-score [a] | total spectral counts (thymidine chase) PRMT1 expressed | total spectral counts (thymidine chase) PRMT1 depleted |
|---|---|---|---|---|---|
| MCM2 | 105 | 92 | -0.42 | 50 | 43 |
| MCM3 | 131 | 111 | -0.49 | 55 | 50 |
| MCM4 | 76 | 71 | -0.23 | 25 | 28 |
| MCM5 | 106 | 93 | -0.39 | 24 | 28 |
| MCM6 | 89 | 96 | 0.19 | 21 | 30 |
| MCM7 | 134 | 117 | -0.37 | 50 | 53 |
| PCNA | 525 | 438 | -0.48 | 99 | 95 |
| RFC1 | 441 | 371 | -0.51 | 18 | 10 |
| RFC2 | 152 | 128 | -0.41 | 3 | 6 |
| RFC3 | 136 | 123 | -0.31 | 0 | 0 |
| RFC4 | 211 | 197 | -0.19 | 9 | 15 |
| RFC5 | 170 | 147 | -0.41 | 0 | 0 |
| ATAD5 | 145 | 87 | -1.24 | 0 | 0 |
| PRIM1 | 41 | 41 | -0.04 | 0 | 0 |
| PRIM2 | 103 | 104 | 0.02 | 0 | 0 |
| POLA1 | 162 | 123 | -0.8 | 0 | 0 |
| POLA2 | 28 | 27 | 0.12 | 0 | 0 |
| POLE | 96 | 82 | -0.51 | 0 | 0 |
| POLD1 | 167 | 154 | -0.25 | 0 | 5 |
| POLD2 | 33 | 28 | -0.53 | 0 | 0 |
| POLD3 | 96 | 80 | -0.55 | 0 | 0 |
| TIMELESS | 11 | 13 | 0.76 | 0 | 0 |
| CLASPIN | 13 | 12 | 0.47 | 0 | 0 |
| WDHD1 | 160 | 119 | -0.73 | 0 | 0 |
| RPA1 | 263 | 240 | -0.29 | 23 | 24 |
| RPA2 | 51 | 58 | 0.38 | 0 | 0 |
| RPA3 | 18 | 18 | 0.26 | 0 | 0 |
| RNASEH2A | 21 | 23 | 0.28 | 0 | 0 |
| RNASEH2B | 71 | 71 | 0.15 | 0 | 0 |
| FEN1 | 244 | 198 | -0.58 | 94 | 91 |
| LIG1 | 148 | 117 | -0.64 | 0 | 0 |
| LIG3 | 116 | 96 | -0.49 | 0 | 0 |
| MSH2 | 610 | 512 | -0.45 | 77 | 97 |
| MSH3 | 313 | 241 | -0.7 | 0 | 0 |
| MSH6 | 789 | 651 | -0.53 | 84 | 75 |
| FANCD2 | 37 | 23 | -1.13 | 0 | 0 |
| FANCI | 124 | 79 | -1.05 | 36 | 31 |
| TOP1 | 312 | 287 | -0.26 | 350 | 338 |
| TOP2A | 560 | 403 | -0.86 | 167 | 144 |
| TOP2B | 271 | 220 | -0.61 | 119 | 123 |
| VCP | 84 | 76 | 0.06 | 65 | 86 |
| RAD18 | 38 | 34 | -0.29 | 0 | 3 |
| RIF1 | 39 | 32 | -0.42 | 0 | 0 |

[a] Ratio of spectral counts in the EdU pulse for PRMT-depleted to spectral counts in the EdU pulse for PRMT1-expressed, averaged across the three replicates, expressed as a z-score.

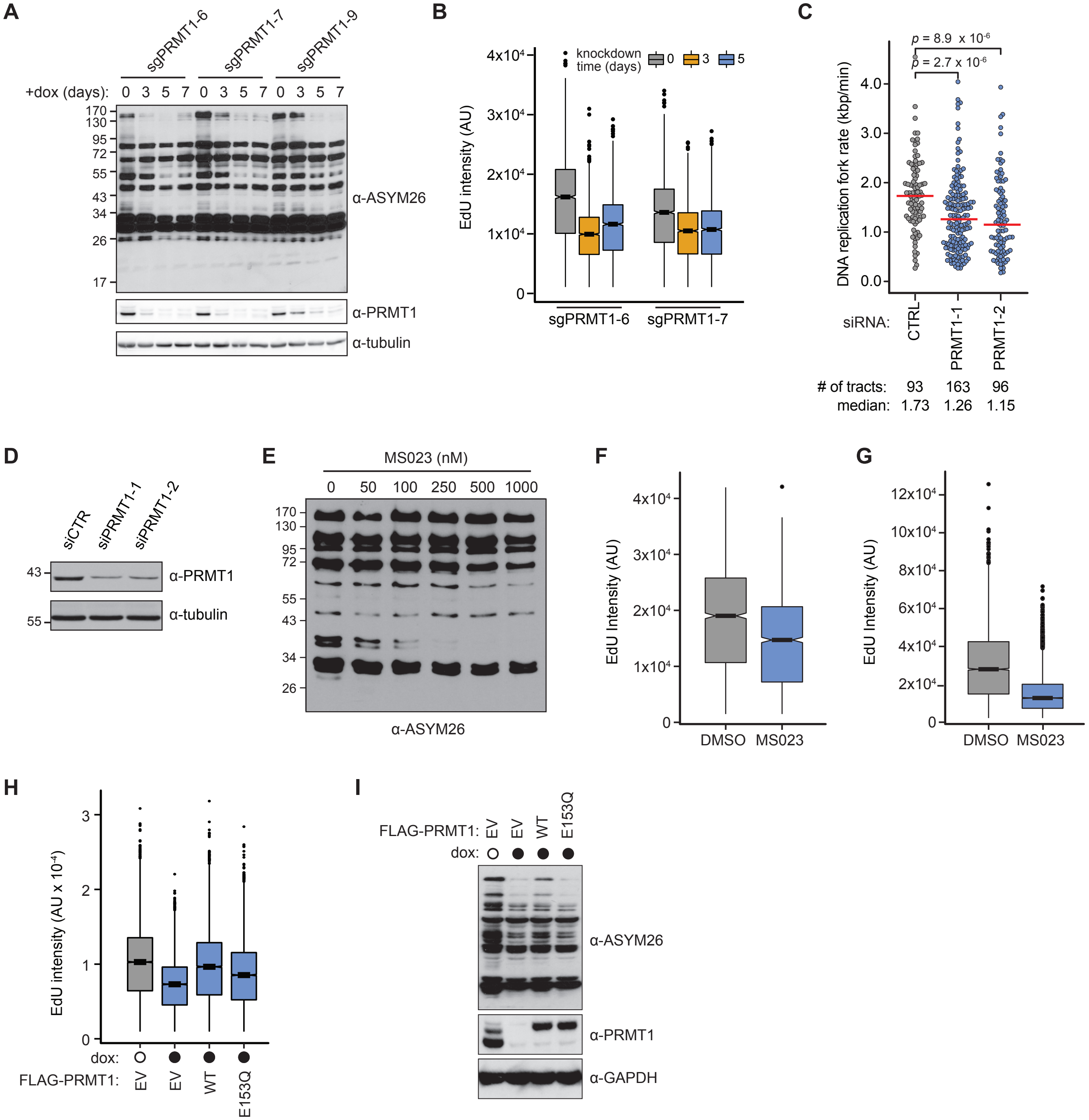

**Figure S1. Relevant to Figure 2.**

**(A)** Global asymmetric arginine dimethylation following *PRMT1* depletion. *PRMT1* CRISPRi cells carrying the indicated sgRNAs were treated with doxycycline and sampled at the indicated times. Whole cell lysates were immunoblotted to detect asymmetric arginine dimethylation ($\alpha$-ASYM26), PRMT1, and tubulin. The positions of molecular weight standards, in kDa, are indicated to the left.

**(B)** EdU incorporation in *PRMT1* CRISPRi cells, replicate two. CRISPRi cells carrying doxycycline-inducible sgRNAs for *PRMT1* were pulse-labelled with EdU at the indicated times after the addition of doxycycline. EdU intensity per cell was measured by flow cytometry and is plotted in arbitrary units as box plots, with horizontal bars indicating the medians. Boxes span the first through third quartiles, whiskers extend to the last data points within 1.5 times the interquartile range, and outliers are plotted as circles. A minimum of 10,000 cells were measured per sample.

**(C)** DNA replication fork rate analysis of U2OS cells following a 2-day siRNA *PRMT1* knockdown. Cells

were sequentially pulse-labelled with CldU and IdU, DNA fibres were isolated and subjected to molecular combing. The IdU tract lengths were measured and are plotted as replication fork rates. Medians are indicated by horizontal red bars. The *p*-values were calculated with a two-sided Mann-Whitney *U* test.
**(D)** PRMT1 protein levels for samples in panel (C) were detected by immunoblotting for PRMT1 and tubulin. The positions of molecular weight standards, in kDa, are indicated to the left.
**(E)** Global asymmetric arginine dimethylation following type I PRMT1 inhibition. HEK293T cells were treated with MS023 at the indicated concentration for 2 days and nuclear extracts were immunoblotted to detect asymmetric arginine dimethylation ($\alpha$-ASYM26), PRMT1, and tubulin. The positions of molecular weight standards, in kDa, are indicated to the left.
**(F)** A375 CRISPRi sgPRMT1-6 cells were treated for 4 days with vehicle (DMSO) or 1 µM MS023, without PRMT1 depletion, followed by pulse-labelling with EdU. The EdU intensity per cell was measured by flow cytometry and is plotted in arbitrary units as box plots, with horizontal bars indicating the medians. Boxes span the first through third quartiles, whiskers extend to the last data points within 1.5 times the interquartile range, and outliers are plotted as circles. A minimum of 6,000 events were analyzed for each condition.
**(G)** U2OS cells were treated for 4 days with vehicle (DMSO) or 1 µM MS023, followed by pulse-labelling with EdU. The EdU intensity per cell was measured by flow cytometry and is plotted in arbitrary units as box plots, with horizontal bars indicating the medians. Boxes span the first through third quartiles, whiskers extend to the last data points within 1.5 times the interquartile range, and outliers are plotted as circles. A minimum of 12,000 events were analyzed for each condition.
**(H)** Rescue of EdU incorporation in *PRMT1* CRISPRi cells, replicate 2. CRISPRi sgPRMT1-7 cells were treated with doxycycline (closed circles) for 5 days to deplete PRMT1, prior to transfection with the empty vector (EV), *FLAG-PRMT1* (WT), or $FLAG\text{-}PRMT1_{E153Q}$ (catalytic-dead; E153Q) prior to pulse-labelling with EdU. The EdU intensity per cell was measured by flow cytometry and is plotted in arbitrary units as box plots, with horizontal bars indicating the medians. Boxes span the first through third quartiles, whiskers extend to the last data points within 1.5 times the interquartile range, and outliers are plotted as circles. A parallel culture without PRMT1 depletion is shown for comparison (open circle).
(I) Whole cell lysates from the samples in (H) were immunoblotted to detect asymmetric arginine dimethylation ($\alpha$-ASYM26), PRMT1, and GAPDH. The positions of molecular weight standards, in kDa, are indicated to the left.

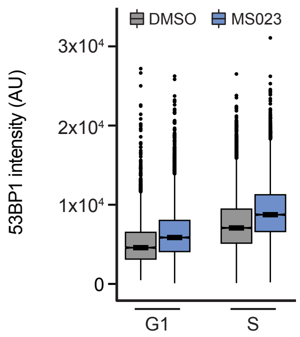

**Figure S2. Relevant to Figure 3.**
**(A)** Quantification of chromatin-bound 53BP1 following PRMT inhibition, replicate 2. U2OS cells were treated with MS023 for 3 days prior to pulse-labelling with EdU. Cells were extracted to remove soluble proteins, stained with DAPI, and subjected to flow cytometry. 53BP1 intensity was measured for each cell in G1 phase or S phase and plotted in arbitrary units as box plots, with horizontal bars indicating the medians. Boxes span the first through third quartiles, whiskers extend to the last data points within 1.5 times the interquartile range, and outliers are plotted as circles. A minimum of 13,000 cells were measured per sample.

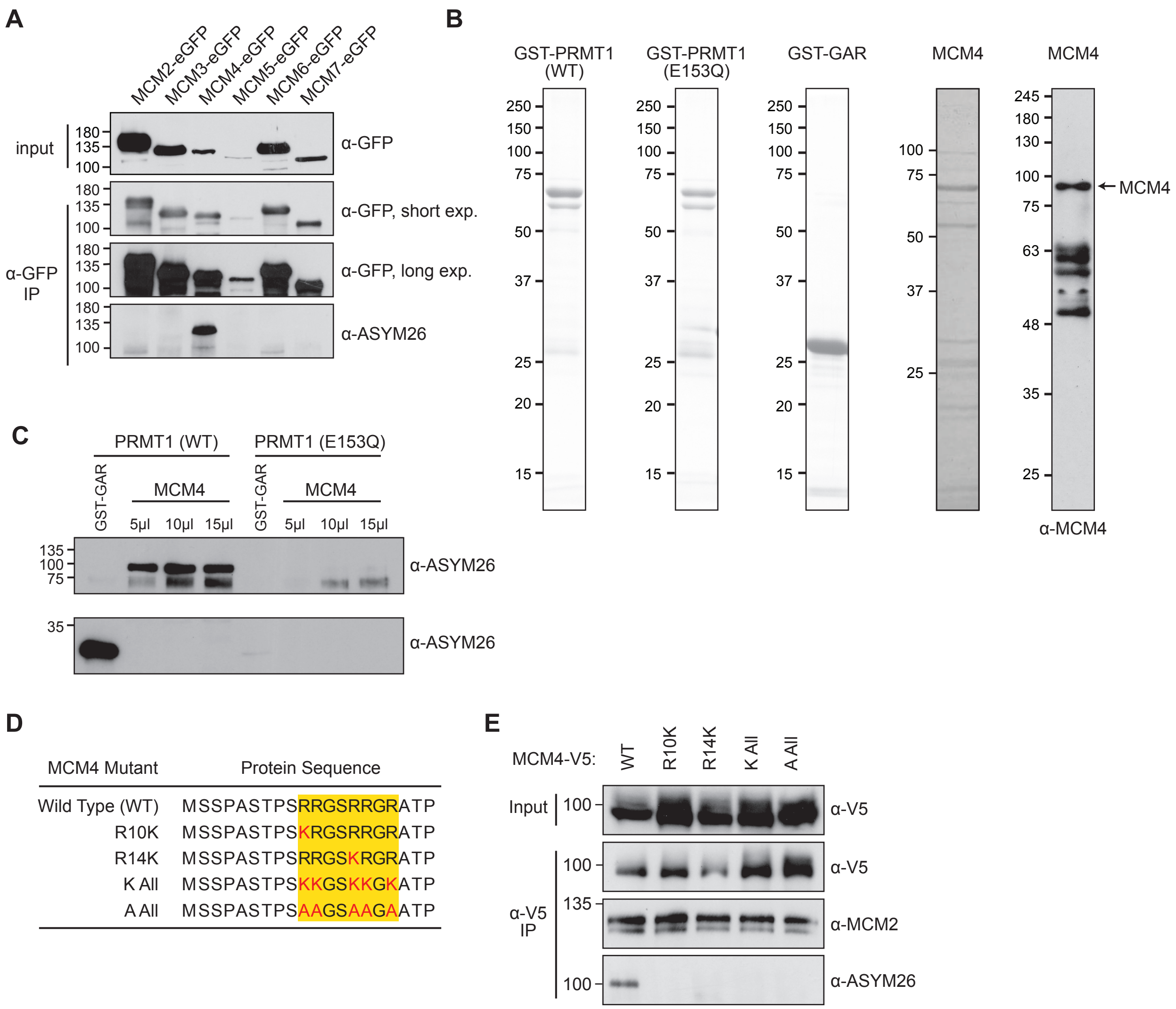

**Figure S3. Relevant to Figure 4.**
**(A)** Asymmetric arginine dimethylation of MCM subunits analyzed by immunoblotting. HEK293T cells were transfected to express the indicated eGFP-tagged MCM proteins, nuclear extracts were prepared (except for eGFP-MCM4 where a cytoplasmic extract was prepared; input), and immunoprecipitated with an anti-GFP antibody (α-GFP IP). Immunoblots were probed to detect the eGFP-tagged proteins (α-GFP) and asymmetric arginine dimethylation (α-ASYM26). The positions of molecular weight standards, in kDa, are indicated to the left.
**(B)** Purified recombinant GST-PRMT1 (WT and E153Q), GST-GAR, and MCM4 were fractionated on SDS-PAGE followed by staining with Coomassie blue. The MCM4 sample was also immunoblotted (far right) and probed with antibody against MCM4 (α-MCM4). The positions of molecular weight standards, in kDa, are indicated to the left.
**(C)** Methylation of recombinant MCM4 with purified GST-PRMT1. MCM4 or GST-GAR was incubated with wild-type (WT) GST-PRMT1, or catalytically inactive (E153Q) GST-PRMT1, for 3 hours at 30°C. The samples were immunoblotted to detect asymmetric arginine dimethylation (α-ASYM26). The positions of molecular weight standards, in kDa, are indicated to the left.
**(D)** Schematic of the MCM4 mutants investigated. The first 20 amino acids of MCM4 are shown. The yellow box highlights the putative GAR motif of MCM4 and the red letters indicate which residues are mutated in each variant.
**(E)** Asymmetric arginine dimethylation of MCM4 variants and interaction with MCM2 analyzed by immunoblotting. HEK293T cells were transfected to express the indicated V5-tagged MCM4 variants, nuclear extracts were prepared (input), and immunoprecipitated with an anti-V5 antibody (α-V5 IP). Immunoblots were probed to detect the V5-tagged MCM4 proteins (α-V5), MCM2 (α-MCM2) and asymmetric arginine dimethylation (α-ASYM26). The positions of molecular weight standards, in kDa, are indicated to the left.

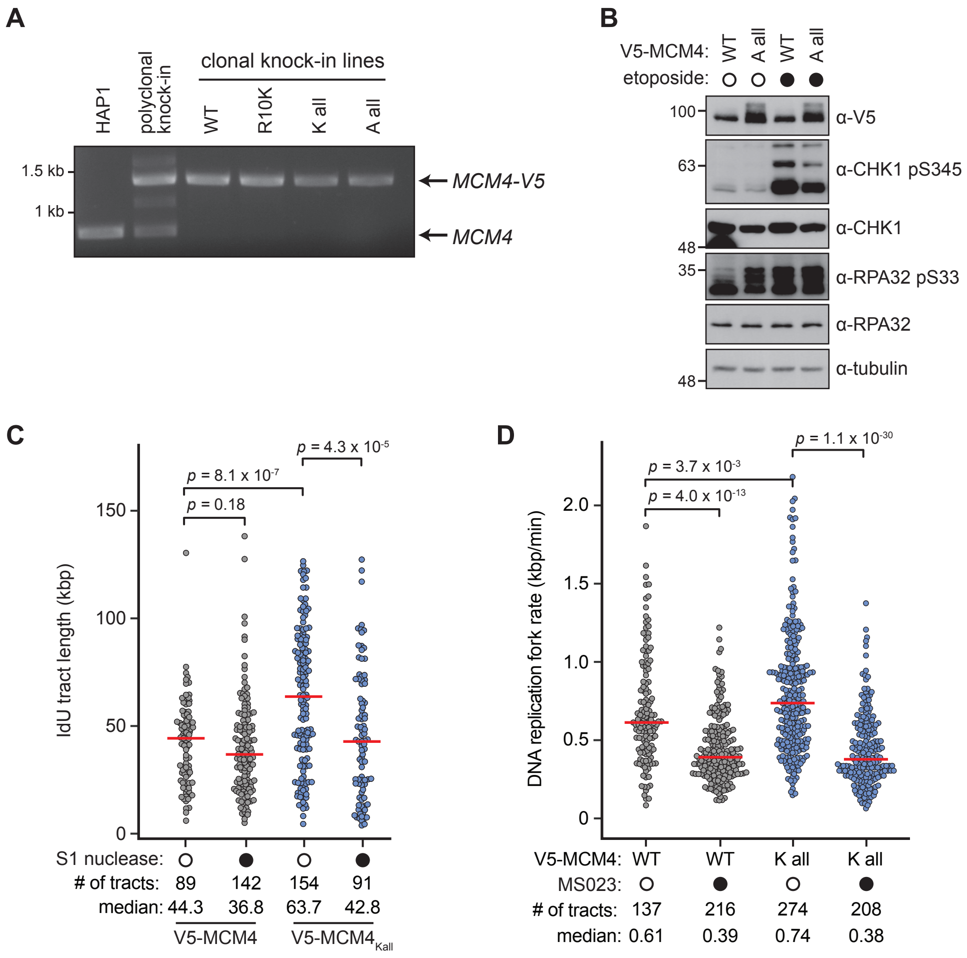

**Figure S4. Relevant to Figure 5.**

**(A)** Genomic DNA from parental HAP1, a representative polyclonal knock-in, or homozygous *MCM4* knock-in lines was PCR-amplified with primers targeting the 5' region of MCM4. The expected positions of the native allele and the V5-tagged knock-in alleles are indicated. The positions of molecular weight standards, in kbp, are indicated to the left.

**(B)** Analysis of CHK1 and RPA32 phosphorylation in *MCM4* variant knock-in cells. Extracts of the indicated knock-in lines were immunoblotted to detect V5-MCM4, phosphorylated CHK1 (CHK1 p345), CHK1, phosphorylated RPA32 (RPA32 pS33), RPA32, and tubulin. Where indicated by the filled circles, cells were treated with 25 μM etoposide for 30 min prior to extraction. The positions of molecular weight standards, in kDa, are indicated to the left.

**(C)** Nuclease sensitivity of replication tracts, replicate two. *MCM4* and *MCM4*$_{Kall}$ knock-in cells were sequentially pulse-labelled with CldU and IdU, followed by DNA fibre preparation, and S1 nuclease treatment as indicated. After molecular combing the IdU tract lengths were measured and are plotted. Medians are indicated by horizontal red bars and *p*-values were calculated with a two-sided Mann-Whitney *U* test.

**(D)** DNA replication fork rate analysis of MCM4 knock-in lines following PRMT inhibition, replicate two. *MCM4* and *MCM4*$_{Kall}$ knock-in cells were treated with 2 μM MS023 for 3 days, followed by sequential pulse-labelling with CldU and IdU, DNA fibre isolation, and molecular combing. The IdU tract lengths were measured and are plotted as replication fork rates. Medians are indicated by horizontal red bars and *p*-values were calculated with a two-sided Mann-Whitney *U* test.

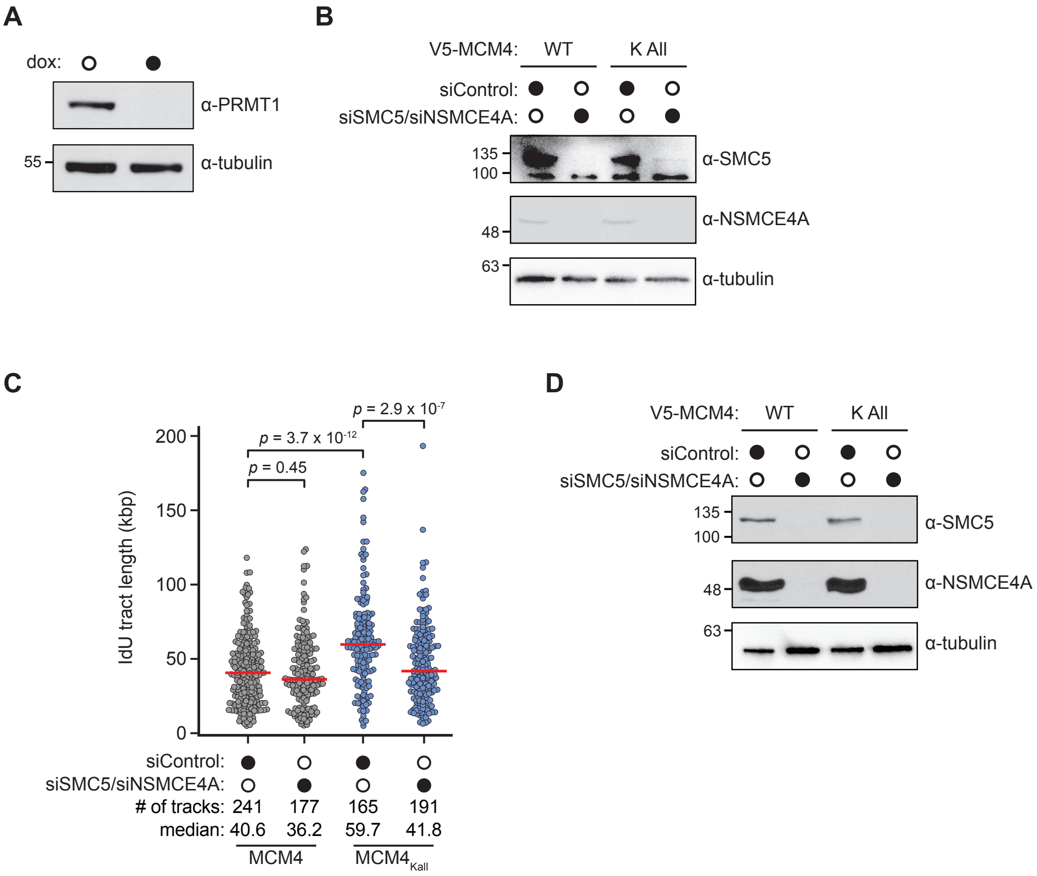

**Figure S5. Relevant to Figure 6.**
**(A)** PRMT1 depletion in a CRISPRi line for differential iPOND. CRISPRi cells carrying a doxycycline-inducible sgRNA for PRMT1 were sampled after 5 days in the presence (closed circle) or absence (open circle) of doxycycline and immunoblotted to detect PRMT1 and tubulin. The positions of molecular weight standards, in kDa, are indicated to the left.
**(B)** Analysis of *SMC5* and *NSMCE4A* siRNA knockdown in *MCM4* variant knock-in cells. Cells were transfected with the indicated (closed circles) siRNAs 3 days prior to harvesting and preparation of extracts. Samples were immunoblotted to detect SMC5, NSMCE4A, and tubulin. Corresponds to Figure 6E.
**(C)** DNA replication tract length analysis of *MCM4* knock-in lines following SMC5/6 complex knockdown, replicate two. *MCM4* and $MCM4_{Kall}$ knock-in cells were treated with siRNAs targeting *NSMCE4A* and *SMC5*, followed by sequential pulse-labelling with CldU and IdU, DNA fibre isolation, and molecular combing. The IdU tract lengths were measured and are plotted. Medians are indicated by horizontal red bars and *p*-values were calculated with a two-sided Mann-Whitney *U* test.
**(D)** Analysis of *SMC5* and *NSMCE4A* siRNA knockdown in *MCM4* variant knock-in cells. Cells were transfected with the indicated siRNAs (closed circles) 3 days prior to harvesting and preparation of extracts. Samples were immunoblotted to detect SMC5, NSMCE4A, and tubulin. Corresponds to panel (C).
